# Neuroanatomy and Behaviour in Mice with a Haploinsufficiency of AT-Rich Interactive Domain 1B (ARID1B) Throughout Development

**DOI:** 10.1101/2020.03.31.017905

**Authors:** J. Ellegood, S.P. Petkova, A. Kinman, L.R. Qiu, A. Wade, D. Fernandes, Z. Lindenmaier, A. Crieghton, L. Nutter, A.S. Nord, J.L. Silverman, J.P. Lerch

## Abstract

One of the causal mechanisms underlying neurodevelopmental disorders (NDDs) is chromatin modification, and genes that regulate chromatin modify and control events regulating the formation of neural connections. AT-Rich Interactive Domain 1B *(ARID1B)*, a chromatin modifier, has been shown to be reduced in autism spectrum disorder (ASD) and to affect rare and inherited genetic variation in a broad set of NDDs. For this work, a novel preclinical mouse model of *Arid1b* deficiency was created and molecularly validated to characterize and define neuroanatomical, behavioural and transcriptional phenotypes. Brains of adult *Arid1b*^*+/-*^ mice had a smaller cerebellum along with a larger hippocampus and corpus callosum. In addition, a notable sex dependence was observed throughout development; males had an early emergence of the neuroanatomical phenotype around postnatal day 7, whereas females had a delayed emergence of the phenotype around postnatal day 40. Behavioural assays relevant to NDD were conducted during neonatal development and adulthood to evaluate general health, anxiety-like, motor, cognitive, and social behaviours in *Arid1b*^+/-^ mice. During neonatal development, *Arid1b*^+/-^ mice exhibited robust impairments in ultrasonic vocalizations (USVs) and metrics of developmental growth. As adults, *Arid1b*^+/-^ mice showed low motor skills in open field exploration and normal three chambered approach. *Arid1b*^+/-^ mice had learning and memory deficits in novel object recognition but surprisingly not in visual discrimination and reversal touchscreen tasks. Social interactions in the male-female social dyad with USVs revealed social deficits on some but not all parameters. No repetitive behaviours were observed. This study represents a full investigation of *Arid1b*^*+/-*^ haploinsufficiency throughout development and highlights the importance of examining both sexes throughout development in NDDs.

## Introduction

Neurodevelopmental disorders (NDDs), including autism spectrum disorder (ASD) and intellectual disability (ID), are prevalent and pervasive lifelong disorders that are behaviourally defined with numerous etiologies and currently no biological markers. The NDD behavioural phenotype is extremely heterogeneous, as are the genetics. For example, the Simons Foundation Autism Research Initiative (SFARI) currently lists more than 900 autism relevant genes (gene.sfari.org)^1^, a number that would increase when all NDDs are taken into account. In an effort to examine the effect that individual genes have on both the brain and behaviour, several groups have started to produce genetically engineered mouse models of the high confidence NDD and ASD relevant genes.

One of the causal mechanisms underlying these disorders is chromatin modification. Genes that regulate chromatin can modify and control events regulating the formation of neural connections; they are also critical for correct epigenetic marking with histone post translational modifications^2^. Chromatin remodeling complexes control the local chromatin state, yielding either transcriptional activation or repression and thus have broad reaching effects on function by regulating numerous gene events. One of these high confidence chromatin-related genes, *ARID1B*, has been shown to be reduced in ASD and affected rare and inherited genetic variation in a broad set of NDD cases^3,4^.

*ARID1B*, and genes in the chromatin modification complex, SWI/SNF, are thought to cause Coffin-Siris syndrome (CSS)^5^. CSS is a neurodevelopmental disorder that is exceedingly rare and clinical presentation includes developmental delay, identified by difficulties with feeding, low birth weight, and general failure to thrive. Other symptoms include ID, impaired motor skills and delayed or absent development of speech and language. Patients present with short stature, broad and coarse facial features, hirsutism, and hypoplasia of the tips of the fingers and toes. Neuroanatomical assessments in the original CSS cases report smaller head circumferences^5^, Dandy Walker malformations (hypoplasia of the cerebellum)^6,7^, and an abnormally thin corpus callosum^8^. In one study, the patient was revealed to have, in addition to a thin corpus callosum, striking deficits in the brainstem and cerebellum. Specifically, ectopic neurons were found throughout the medulla, inferior olive, and the white matter of the cerebellum^8^. In a recent report of 143 different patients with *ARID1B* mutations, these original observations about CSS remain consistent, with agenesis or hypoplasia of the corpus callosum prevalent in 43% of patients^9^.

Neuroanatomical assessments in the brains of mouse models of NDD have become more frequent with full brain phenotypes being established and characterized in several different rodent models^10-16^. However, with well over nine hundred genes linked to NDD, characterization of the more prominent and closely linked genetic models, like *Arid1b*, are imperative. Furthermore, characterizing the functional impact of a germline *Arid1b* mutation on brain development, growth, and behaviour *in vivo* could reveal specific and generalizable mechanisms linking chromatin biology to pathology and outcomes. Moreover, having a rigorous reproducible model system allows for the testing of medicinal or genetic interventions and developing therapeutics. Three groups have recently published papers examining mouse models of *Arid1b* haploinsufficiency^17-19^. In those studies, the authors report several behavioural abnormalities relevant to ASD phenotypes, including cognitive and social behaviour deficits. Neuroanatomical volumetric differences were also found in the dentate gyrus and corpus callosum that were consistent with the known neuroanatomy in a subset of human patients with CSS. However, in none of these previous investigations of *Arid1b* mutant mice was a full brain neuroanatomical assessment performed. One study focused on only two areas relevant to CSS^17^, and the other two groups did not specifically look for neuroanatomical alterations^18,19^, but rather focussed on smaller microscopic differences in cortical structures. Moreover, these studies examined limited windows of time, focusing only on adulthood for their neuroanatomical and behavioural assessments.

To better understand both the behaviour and the neuroanatomical phenotype throughout development, we generated a novel preclinical model of *Arid1b* using CRISPR/Cas9 genome engineering. We assayed neuroanatomical, behavioural and transcriptional phenotypes associated with *Arid1b* haploinsufficiency in the mouse. Behavioural assays were conducted during both neonatal development and adulthood. Additionally, using *in vivo* manganese enhanced MRI (MEMRI), a developmental time course of the neuroanatomy was assessed throughout the entire brain from birth into adulthood.

## Methods

### Generation of the Mice

Precision genetic engineering via the Cas9/CRISPR system was used to generate one of the first *Arid1b* mutant mouse lines. This allele from project TCPR0317 was generated at The Centre for Phenogenomics (TCP) by injecting Cas9 nickase (D10A) and single guide RNAs with spacer sequences of CTGCTTAGCAAGTTACCACT and GCCTGATACAGCACTTACAT targeting the 5’ side and ACACTAAAGGGGTTGCTTTC and CTTGTAATCCCCCTGTAGTA targeting the 3’ side of exon OTTMUSE00000314956 (exon 5) resulting in deletion of Chr17 from 5242523 to 5243410 with insertion of TT. Consistent with earlier studies, this mutation is associated with early embryonic lethality in homozygous carriers. The mouseline C57Bl/6N-*Arid1b*^em1(IMPC)Tcp^ was made as part of the NorCOMM2 project funded by Genome Canada and the Ontario Genomics Institute at The Centre for Phenogenomics. It was obtained from the Canadian Mutant Mouse Repository (CMMR).

### RNA-Sequencing

Transcriptomic analysis was performed as before^10^. Cerebellar tissue was micro-dissected from adult heterozygous *Arid1b* mice and Wild-Type (WT) littermates (postnatal day (PND)140-150). Stranded mRNA sequencing libraries were prepared using a TruSeq Stranded mRNA kit with all four samples pooled and sequenced in one lane on the Illumina HiSeq platform using a single-end 100-bp strategy. Each library was quantified and pooled before sequencing at the UC Davis DNA Technologies Core. Reads from RNA-seq were aligned to the mouse genome (mm9) using STAR (version 2.5.3a). Aligned reads mapping to genes were counted at the gene level using subreads featureCounts. The mm9 knownGene annotation track and aligned reads were used to generate quality control information using the full RSeQC tool suite. Unaligned reads were quality checked using FastQC.

### Differential Expression Analysis

Raw count data for all samples were used for differential expression analysis using edgeR. Genes with at least 20 reads per million in at least one sample were included for analysis, resulting in a final set of 8,942 genes for differential testing. Tagwise dispersion estimates were generated and differential expression analysis was performed using a generalized linear model with genotype as the variable for testing. Normalized expression levels were generated using the edgeR rpkm function. Normalized log2(RPKM) values were used to plot expression data for *Arid1b*. Pseudo-count values were used to plot the summary heatmap. Mouse gene ontology (GO) data was downloaded from Bioconductor (org.Mm.eg.db). We used the goseq package to test for enrichment of GO terms indicating parent:child relationships. For GO analysis, we examined down- and up-regulated genes separately for genes meeting an FDR < 0.2. For the enrichment analysis, the test set of differentially expressed genes was compared against the background set of genes expressed in our study based on minimum read-count cutoffs described above.

### Breeding

All animals were housed in a temperature-controlled vivarium maintained on a 12-hour light-dark cycle. All procedures were approved by the Institutional Animal Care and Use Committee either at The Centre for Phenogenomics (TCP) in Toronto, Ontario, Canada, or at the University of California Davis School of Medicine in Sacramento, CA, USA, and were conducted in accordance with the Canadian Council on Animal Care Guide to Care and Use of Experimental Animals or the National Institutes of Health Guide for the Care and Use of Laboratory Animals. Efforts were made to minimize animal suffering and the number of animals used.

### Perfusions

In total, 47 fixed mouse brains were examined: 14 *Arid1b*^*+/-*^ mice (7 females, 7 males) and 33 WT (C57Bl6/NCrl, 13 females and 20 males). All mice were perfused at PND60. Mice were anesthetized with a mixture of ketamine/xylazine and intracardially perfused with 30mL of 0.1M PBS containing 0.05 U/mL heparin (Sigma) and 2mM ProHance (Bracco Diagnostics, a Gadolinium contrast agent) followed by 30 mL of 4% paraformaldehyde (PFA) containing 2mM ProHance^20,21^. Perfusions were performed with a minipump at a rate of approximately 1 mL/min. After perfusion, mice were decapitated and the skin, lower jaw, ears, and the cartilaginous nose tip were removed. The brain and remaining skull structures were incubated in 4% PFA + 2mM ProHance overnight at 4°C then transferred to 0.1M PBS containing 2mM ProHance and 0.02% sodium azide for at least 1 month prior to MRI scanning ^22^.

### Magnetic Resonance Imaging – Ex Vivo

A multi-channel 7.0 Tesla MRI scanner (Agilent Inc., Palo Alto, CA) was used to image the brains within their skulls. Sixteen custom-built solenoid coils were used to image the brains in parallel^21,23^. In order to detect volumetric changes, we used the following parameters for the MRI scan: T2-weighted, 3-D fast spin-echo sequence, with a cylindrical acquisition of k-space, a TR of 350 ms, and TEs of 12 ms per echo for 6 echoes, field-of-view equaled to 20 x 20 × 25 mm^3^ and matrix size equaled to 504 × 504 × 630. Our parameters output an image with 0.040 mm isotropic voxels^24^.The total imaging time was 14 hours.

### Mice for Longitudinal MRI

Mice were bred by crossing *Arid1b*^*+/-*^ with a C57Bl6/NCrl mice to produce heterozygous offspring. The number of mouse pups in each litter was reduced to 6 to ensure equal manganese intake by pups through maternal milk. On PND2, mice were tattooed with green ink on their paws for identification and tail clipped for genotyping. Male and female mice were scanned longitudinally in two cohorts over 8 different time-points: PND4, 6, 8, 10, 17, 23, 35, and 60. Cohort 1 was scanned over the first five timepoints, and cohort two was scanned over all 8 timepoints to extend our developmental window to PND60. Cohort 1 consisted of 11 males and 11 females, 5 of which were mutants and 6 of which were WT controls of each sex. Cohort 2 consisted of 18 males and 18 females, 9 of which were mutants and 9 of which were WT controls of each sex. For the first five developmental timepoints, group sizes ranged from 12-19 mice per sex per genotype; for the final 3 timepoints group sizes ranged from 7-9 mice per sex per genotype. An incomplete group at a given timepoint was due to a loss of either the scan (corruption, movement etc.) or the animal itself during the longitudinal time-course.

### Magnetic Resonance Imaging – In Vivo

Findings in the *ex vivo* experiments, and the fact that *Arid1b* haploinsufficiency is relevant to NDDs, highlighted the need to examine the developmental time course of the neuroanatomical differences. Therefore, a longitudinal study was performed to track the development of the neuroanatomy from birth to adulthood, similar to previous work^25,26^. 24 hours prior to the scan, a 0.4 mmol/kg dose of 30mM manganese chloride (MnCl_2_) was administered as a contrast agent. For mice that were 10 days old or younger, the dam was intraperitoneally injected with MnCl_2_, and pups received MnCl_2_ through maternal milk. Mice that were older than ten days received intraperitoneal injections of MnCl_2_. Up to four mice were scanned simultaneously. Custom built 3D printed holders, which allowed for anesthetic delivery and scavenging, as well as heating were used. During the scan, mice were anesthetized with 1-2% isoflurane, and respiratory rate was monitored using a self-gated signal from a modified 3D gradient echo sequence ^27^.

A multi-channel 7.0 Tesla MRI scanner with a 30 cm diameter bore (Bruker, Billerica, Massachusetts), equipped with 4 individual cryogenically cooled coils was used to acquire images of the mouse brains. Parameters of the scan are as follows: T1 weighted FLASH 3D gradient echo sequence, TR = 26 ms, TE = 8.250 ms, flip angle = 23°, field of view = 25 × 22 × 22 mm, with a matrix size of 334 × 294 × 294 yielding an isotropic imaging resolution of 75 μm. Imaging time was 58 minutes. After imaging, mice were transferred to a heated cage for 5-10 minutes to recover from the anesthesia, and then returned to their home cage.

### Imaging Registration and Analysis

Image registration is necessary for quantifying the anatomical differences across images. For both the *in vivo* and *ex vivo* images the registration consisted of both linear (rigid then affne) transformations and non-linear transformations. These registrations were performed with a combination of mni_autoreg tools^28^ and ANTS (advanced normalization tools)^29,30^. A two-level registration pipeline was used on the *in vivo* images and has been detailed in previous studies^25^. Briefly, images from the same age were registered together in the first level creating a consensus average for the brain at each age. For example, all of the PND4 images, both mutant and WT, were registered together to create a PND4 average brain, all of the PND6 images are registered together to create a PND6 average brain, etc. In the second level, each age-consensus average was registered to the average of the next-oldest age (I.e. PND4 average registered to the PND6 average, PND6 to the PND8, etc.) Transformations from each level were then concatenated so that every single image in the longitudinal dataset could be mapped to the final PND60 average. For the *ex vivo* image registration, a simpler one-level registration pipeline is used. After registration, all scans were resampled with the appropriate transform and averaged to create a population atlas representing the average anatomy of the study sample. The result of these registrations were deformation fields that transform images to a consensus average. Therefore, these deformations fields quantify anatomical differences between images.

As detailed in previous studies^31,32^, the Jacobian determinants of the deformation fields were computed and analyzed to measure the volume differences between subjects at every voxel. A pre-existing classified MRI atlas was warped onto the population atlas (containing 282 different segmented structures encompassing cortical lobes, large white matter structures such as the corpus callosum, ventricles, cerebellum, brain stem, and olfactory bulbs^25,33-36^) to compute the volume of brain structures in all the input images. A linear model with a genotype predictor was used to assess the significance of genotype. The model was either fit to the volume of every structure independently (structure-wise statistics) of fit to every voxel independently (voxel-wise statistics), and multiple comparisons in this study were controlled for using the False Discovery Rate^37^.

For the longitudinal analysis, a linear mixed effects model was fit to the volume of every structure with fixed effects of sex, genotype, age, sex-genotype interaction, sex-age interaction, genotype-age interaction, sex-genotype-age interaction, and whole-brain volume; and a random intercept for each mouse. To assess the significance of an effect, a reduced model (model without the particular predictor) was fit to the data and the log likelihood ratio test was used to assess whether the particular effect was significantly important in predicting the data. For example, to evaluate the genotype effect, the reduced model contained fixed effects of sex, age, and sex-age interaction, and significant log likelihood ratio test for a structure results indicate genotype had a notable effect on the volume of that structure. To estimate direction of effect, Satterthwaite approximation^38^ was used to estimate degrees-of-freedom.

### Behavioural Testing

Behavioural experiments were performed on separate cohorts of mice at UC Davis. Cohort 1 was tested through either ultrasonic vocalizations or postnatal developmental physical milestones and neurological reflexes PND2-14, and then they were again tested through adult behaviours consisting of elevated plus maze, light↔dark exploration, open field arena exploration, novel object recognition, self-grooming, three chambered social approach, male female dyad social interactions, fear conditioning, hot plate and grip strength starting at 10 weeks of age. Cohort 2 was tested through either ultrasonic vocalizations or developmental milestones PND2-14 and were tested through touchscreen starting at 8 weeks of age. Lethal dose of PTZ served as the endpoint for both cohorts at 18 weeks of age. To minimize the carry-over effects from repeated testing, assays were performed in the order of least to the most stressful tasks. Cohort 1 was sampled from 12 litters and was tested in elevated plus maze, light↔dark exploration, novel object recognition, social approach, male female social dyad interactions, self groom, contextual and cued fear conditioning, hot plate, grip strength and lethal dose PTZ. Cohort 2 was sampled from 12 litters and was tested in pairwise discrimination and reversal touchscreen task and lethal dose of PTZ. All male and female mice were used from every litter. In cohort 1, the order of testing was as follows with at least 48-hrs separating tasks: (1) PND3, 5, 7, 9 ultrasonic vocalizations or PND2, 4, 6, 8, 10, 12, 14 developmental milestones, (2) elevated plus-maze at 10 weeks of age, (3) light↔dark exploration task at 10 weeks of age, (4) open field locomotion at 11 weeks of age, (5) balance beam walking at 11 weeks of age, (6) spontaneous alternation at 12 weeks of age, (7) rotarod at 12 weeks of age, (8) novel object recognition at 13 weeks of age, (9) self-groom at 15 weeks of age, (10) social approach at 15 weeks of age, (11) male female social dyad interaction at 16 weeks of age, (12) grip strength at 17 weeks of age, (13) hot plate at 17 weeks of age, (14) fear conditioning at 18 weeks of age, and (15) lethal dose chemoconvulsant at 19 weeks of age. In cohort 2, ultrasonic vocalizations or developmental milestones were conducted between PND2 and 14 in separate litters, weight restriction for touchscreen assay began at 7 weeks of age, pretraining began at 9 weeks of age and touchscreen testing continued daily until completion, DigiGait was conducted at 17 weeks of age and lethal dose chemoconvulsant at 18 weeks of age. All behavioural assays in all cohorts were performed between 9 am-4 pm PST (ZT2-ZT9).

### Developmental Milestones

Developmental milestones were measured on PND2-14, similar to those previously described^39,40^ but elaborated based on the unique features of this line. All measures were conducted by an experimenter blind to genotype and sex. Pups were separated from the dam for a maximum time of four minutes. Pups were observed for abnormalities in eye, pinnae, whisker, and fur development each testing day. Pups were also picked up to observe limb movement for early signs of hypotonia. Physical feature measurements such as body weight and length (subtraction of total length from tip of nose to base of tail and tail length), and head width was measured using a scale (grams) and a digital caliper (mm). Developmental motor milestones to detect neonatal hypotonia and ataxia included negative geotaxis, cliff avoidance, righting reflex and circle transverse. Negative geotaxis was tested by placing each pup, facing downwards, on a surface angled at 45° from parallel, and measuring the time for it to completely turn 180° and face up the incline. Failures to turn and climb were recorded as a maximum score of 30-s. Cliff avoidance was tested by gently placing each pup near the edge of a cardboard box and measuring the time for it to turn its head or back away from the edge. Failures to avoid the cliff was recorded as a maximum score of 30-s. Righting reflex was tested by placing each pup on its back, releasing it, and measuring the time for it to fully flip over onto four paws on each developmental day. Righting reflex, negative geotaxis and cliff aversion all test early limb coordination, vestibular system and proprioception. Circle transverse was tested by placing each pup in the center of a circle with a 5″ (12.5 cm) diameter drawn on a laminated sheet of 8.5″×11″ white paper and measuring the time for it to exit the circle. Circle transverse is an early metric of motor ability and ambulation; developmental delay in crawling and walking may be observed using this task. Failures to exit the circle were recorded as a maximum score of 30-s. Milestones for tracking development of fore and hind limb strength were forelimb grasp, forelimb bar holding, forelimb hang, and hindlimb hang. Forelimb grasp tests a mouse’s ability to curl its paw towards an object, a thin wooden stick. Pups typically develop this ability by PND5 and day of first presentation is noted for each pup. Starting around PND6, pups develop enough limb strength to hold themselves up by the forelimbs. Pups are placed by the forelimbs on an unfolded paper clip and time spent able to hang on independently, up to 30 seconds, is recorded. Hindlimb strength testing can be performed earlier than forelimb strength. Once pups develop some limb strength and the grasp reflex, a thin unfolded paper clip bar is placed into their paws and time able to independently hold the bar, up to 30 seconds, is recorded. This task requires high compliance and tested starting at PND8. The delay in testing from PND2 to PND8 is the result of this skill’s difficulty to assess. Subjects must grasp, comply for a set amount of time and hold the bar. Hindlimb hang is performed by placing the pup on the thin edge of a surface or container by its legs and allowing the pup to hang down and support itself on the edge by its hindlimbs. Time spent independently hanging by its hindlimbs, up to 30 seconds, is recorded. Strength tests allow for the determination if motor deficits are due to peripheral limb weakness or centralized neural coordination dysfunction.

### Ultrasonic Vocalizations

Pups were removed individually from the nest at random and gently placed into an isolation container (8 cm × 6 cm × 5 cm; open surface) made of glass. The isolation container was surrounded by a sound attenuating box (18 cm × 18 cm × 18 cm) made of 4 cm thick styrofoam walls. Ultrasonic vocalizations were monitored by an UltraSound Gate Condenser Microphone CM 16 (Avisoft Bioacoustics, Berlin, Germany) attached to the roof of the sound attenuating box, 7 cm above the floor. The microphone was connected via an UltraSoundGate 116 USB audio device (Avisoft Bioacoustics) to a personal computer where acoustic data was recorded with a sampling rate of 250,000 Hz by Avisoft RECORDER (version 2.97; Avisoft Bioacoustics). At the end of the 3-min recording session, both body weight and body temperature were collected. The isolation container was cleaned with 70% ethanol before the beginning of the first test session and between each new litter. Ultrasonic vocalization spectrograms were displayed using Avisoft software and calls were identified manually by a highly trained investigator blinded to genotype.

### Elevated Plus-Maze

Conflict anxiety-like behaviour via the elevated plus maze was measured according to previously described procedures ^41,42^ using a standard mouse apparatus (Med Associates, St. Albans, VT). The maze had two open arms (35.5 cm × 6 cm) and two closed arms (35.5 cm × 6 cm) radiating from a central area (6 cm × 6 cm). A 0.5 cm high lip surrounded the edges of the open arms. 20 cm high walls enclosed the closed arms. The apparatus was cleaned with 70% ethanol before the beginning of the first test session and after each subject mouse. Room illumination was ∼ 300 lux.

### Light ↔ Dark Transitions

Light ↔ dark exploration was measured according to previously published procedures^41,42^. Subject mice were placed in the brightly illuminated, large chamber. The smaller dark chamber (∼ 5 lux) was entered by traversing a partition between the two chambers. Subject mice freely explored for 10 minutes. Time in the dark side chamber and total number of transitions between the light and dark side chambers were automatically recorded using Labview 8.5.1 software (National Instruments, Austin, TX). Room illumination was ∼ 400 lux.

### Open Field

General exploratory locomotion in a novel open field environment was assayed as previously described^41,42^. Open field activity was considered an essential control for effects on physical activity, e.g. sedation or hyperactivity, which could confound the interpretation of results from the reciprocal interactions, self-grooming, novel object recognition, fear conditioning and social approach tasks^43,44^. The testing room was illuminated at ∼30 lux.

### Gait

Treadmill gait analysis was performed using the DigiGait™ system (Mouse Specifics Inc., USA)^45^. Mouse paws were painted with non-toxic red food coloring to augment dark green paw tattoos that generated conflict in DigiGait™ analysis one minute prior to introduction to the walking chamber to reliably capture the entire paw surface area. Before data collection, each subject was acclimated to the Perspex walking chamber for one minute and the treadmill was slowly accelerated to the final speed of 20cm/s to allow mice to adjust to walking on the belt. Digital images of paw placement were recorded through a clear treadmill from the ventral plane of the animal. Mice were tested in a single session at a 20 cm/s treadmill speed maintaining a normal pace walk for WT mice. Non-performers were defined as mice who were unable to sustain walking at 20 cm/s without colliding with the posterior bumper for at least three seconds. There is no practice effect or repeated exposure, and therefore, mice were allowed retrial and retest if they were unable to adjust to walking on the belt easily. The treadmill belt and the encasing Perspex chamber were cleaned with 70% (v/v) ethanol in between tests. For each mouse, videos of ∼5 second duration of all sessions were analyzed using the DigiGait™ Imaging and Analysis software v12.2 (Mouse Specifics Inc., USA). Contrast filters were determined on a mouse-by-mouse case to facilitate consistent recognition of all four paws. All analysis was conducted in a single session by experimenter blind to genotype. Stride length (distance a paw makes during a single stride) and frequency (number of strides per second to maintain pace) were automatically calculated. Data was averaged between left and right paws for fore and hind paws.

### Novel Object Recognition

The novel object recognition (NOR) test was conducted in opaque matte white (P95 White, Tap Plastics, Sacramento, CA) open field arenas (41 cm × 41 cm × 30 cm), using methods similar to those previously described^10,40,42^. The experiment consisted of four sessions: a 30-min exposure to the open field arena the day before the test, a 10-min re-habituation on test day, a 10-min familiarization session and a 5-min recognition test. On day 1, each subject was habituated to a clean, empty open field arena for 30-min. 24-hr later, each subject was returned to the open field arena for another 10-min for the habituation phase. The mouse was then removed from the open field and placed in a clean temporary holding cage for approximately 2-min. Two identical objects were placed in the arena. Each subject was returned to the open field in which it had been habituated and allowed to freely explore for 10-min. After the familiarization session, subjects were returned to their holding cages, which were transferred from the testing room to a nearby holding area. The open field was cleaned with 70% ethanol and let dry. One clean familiar object and one clean novel object were placed in the arena, where the two identical objects had been located during in the familiarization phase. 60-min after the end of the familiarization session, each subject was returned to the arena for a 5-min recognition test, during which time it could to freely explore the familiar object and the novel object. The familiarization session and the recognition test were recorded and scored with Ethovision XT videotracking software (Version 9.0, Noldus Information Technologies, Leesburg, VA). Object investigation was defined as time spent sniffing the object when the nose was oriented toward the object and the nose–object distance was 2-cm or less. Recognition memory was defined as spending significantly more time sniffing the novel object compared to the familiar object via a Student’s paired t-test. Total time spent sniffing both objects was used as a measure of general exploration. Time spent sniffing two identical objects during the familiarization phase confirmed the lack of an innate side bias. Objects used were plastic toys: a small soft plastic orange safety cone and a hard, plastic magnetic cone with ribbed sides, as previously described ^46^.

### Repetitive Self-Grooming

Spontaneous repetitive self-grooming behaviour was scored as previously described^47,48^. Each mouse was placed individually into a standard mouse cage, (46 cm length × 23.5 cm wide × 20 cm high). Cages were empty to eliminate digging in the bedding, which is a potentially competing behaviour. The room was illuminated at ∼40 lux. A front-mounted CCTV camera (Security Cameras Direct) was placed at ∼1m from the cages to record the sessions. Sessions were video-taped for 20 min. The first 10-min period was habituation and was unscored. Each subject was scored for cumulative time spent grooming all the body regions during the second 10 min of the test session. Scoring was conducted by an experimenter blind to genotype.

### Social Approach

Social approach was tested in an automated three-chambered apparatus using methods similar to those previously described ^49-51^. Automated Ethovision XT videotracking software (Version 9.0, Noldus Information Technologies, Leesburg, VA) and modified non-reflective materials for the chambers were employed to maximize throughput. The updated apparatus (40cm x 60cm x 23cm) was a rectangular, three-chambered box made from matte white finished acrylic (P95 White, Tap Plastics, Sacramento, CA). Opaque retractable doors (12cm x 33cm) were designed to create optimal entryways between chambers (5cm x 10cm), while providing maximal manual division of compartments. Three zones, defined using the EthoVision XT software, detected time in each chamber for each phase of the assay. Zones were defined as the annulus extending 2 cm from each novel object or novel mouse enclosure (inverted wire cup, Galaxy Cup, Kitchen Plus, http://www.kitchen-plus.com). Direction of the head, facing toward the cup enclosure, defined sniff time. A top mounted infrared sensitive camera (Ikegami ICD-49, B&H Photo, New York, NY) was positioned directly above every two 3-chambered units. Infrared lighting (Nightvisionexperts.com) provided uniform, low level illumination. The subject mouse was first contained in the center chamber for 10 minutes, then explored all three empty chambers during a 10-minute habituation session, then explored the three chambers containing a novel object in one side chamber and a novel mouse in the other side chamber. Lack of innate side preference was confirmed during the initial 10 min of habituation to the entire arena. Novel stimulus mice were 129Sv/ImJ, a relatively inactive strain, aged 10–14 weeks old, and matched to the subject mice by sex. Number of entries into the side chambers served as a within-task control for levels of general exploratory locomotion.

### Male Female Social Dyad Interaction

The male-female social dyad interaction test was conducted as previously described ^42,48,50^. Briefly, each freely moving male subject was paired for 5-min with a freely moving unfamiliar estrous WT female. A closed-circuit television camera (Panasonic, Secaucus, NJ, USA) was positioned at an angle from the Noldus PhenoTyper arena (Noldus, Leesburg, VA) for optimal video quality. An ultrasonic microphone (Avisoft UltraSoundGate condenser microphone capsule CM15; Avisoft Bioacoustics, Berlin, Germany) was mounted 20 cm above the cage. Sampling frequency for the microphone was 250 kHz, and the resolution was 16 bits. The entire apparatus was contained in a sound-attenuating environmental chamber (Lafayette Instruments, Lafayette, IN) under red light illumination (∼ 10 lux). Duration of nose-to-nose sniffing, nose-to-anogenital sniffing, and following were scored using Noldus Observer 8.0XT event recording software (Noldus, Leesburg, VA) as previously described^52^. Ultrasonic vocalization spectrograms were displayed using Avisoft software and calls were identified manually by a highly trained investigator blinded to genotype.

### Fear conditioning

Delay contextual and cued fear conditioning was conducted using an automated fear-conditioning chamber (Med Associates, St Albans, VT, USA) as previously described ^42^. The conditioning chamber (32 × 25 × 23 cm^3^, Med Associates) was interfaced to a PC installed with VideoFreeze software (version 1.12.0.0, Med Associates) and enclosed in a sound-attenuating cubicle. Training consisted of a 2-min acclimation period followed by three tone-shock (CS–US) pairings (80 dB tone, duration 30 s; 0.5 mA footshock, duration 1 s; intershock interval 90 s) and a 2.5-min period, during which no stimuli were presented. The environment was well lit (∼100 lux), with a stainless-steel grid floor and swabbed with vanilla odor cue (prepared from vanilla extract; McCormick; 1:100 dilution). A 5-min test of contextual fear conditioning was performed 24 h after training, in the absence of the tone and footshock, but in the presence of 100 lux overhead lighting, vanilla odor and chamber cues identical to those used on the training day. Cued fear conditioning, conducted 48 h after training, was assessed in a novel environment with distinct visual, tactile and olfactory cues. Overhead lighting was turned off. The cued test consisted of a 3-min acclimation period followed by a 3-min presentation of the tone CS and a 90-s exploration period. Cumulative time spent freezing in each condition was quantified by VideoFreeze software (Med Associates).

### Touchscreen pairwise discrimination

Pairwise visual discrimination was tested in the automated Bussey-Saksida touchscreen apparatus for mice (Campden Instruments Ltd/Lafayette Instruments, Lafayette, IL, USA), using a procedure modified from original methods described previously ^51,53-55^. The reinforcer was 20 ul of a palatable liquid nutritional supplement (Strawberry Ensure Plus, Abbott, IL, USA) diluted to 50% with water. Each session was conducted under overhead lighting (∼60 lux). A standard tone cue was used to signal the delivery of the reinforcer during pre-training and acquisition. Prior to pre-training, subject mice were weighed, and placed on a restricted diet of 2–4 g of rodent chow per mouse per day, to induce a 15% weight loss. Body weight was carefully monitored throughout the experiment, to ensure that a minimum of 85% of free feeding body weight was maintained for each mouse. The pre-training consisted of four stages as previously published^51^. Stage 1 consisted of two days of habituation (20 min on day 1, and 40 min on day 2) to the chamber and the liquid diet with no images on the screen under overhead lighting (∼60 lux). Stage 2 was a single 45-min session in which entering and exiting the food magazine initiates the next trial and triggers additional reward under overhead lighting. During Stage 3, subjects were trained in daily 45-min sessions during which an image (a random picture from a selection of 40 images) was presented in one of the two windows and remained on the screen until it was touched. Mice must complete 30 trials/day for two consecutive days in order to advance to the next stage. In Stage 4, subjects were trained in 45-min daily sessions in which touching the blank side of the screen was discouraged with a 5-s time-out during which the overhead lighting turned off. Completion of at least 30 trials, at an average accuracy of 80%, on two consecutive days, is required for advancement. Images used in Stages 3 and 4 were not used in the subsequent discrimination task. Only mice that completed all stages of pre-training were advanced to the pairwise visual discrimination task. Subjects were trained to discriminate between two novel images, a spider and an airplane, presented in a spatially pseudo-randomized manner in the two windows of the touchscreen. Each 45-min session consisted of unlimited number of trials separated by 15-s inter-trial intervals (ITI). Designation of the correct and incorrect images was counterbalanced across mice within each genotype. Correct responses were rewarded. Each incorrect response was followed by a correction trial in which the images were presented in an identical manner to the previous trial, until a correct response was made. Criterion was completing at least 30 trials, at an accuracy of 80% or higher, on two consecutive days. Days to reach criterion and percentage of mice reaching criterion on each day were compared between genotypes. After successful completion of the task, mice underwent task reversal in which the opposite image was now correct. Criterion was completing at least 30 trials, at an accuracy of 75% or higher, on two consecutive days. Days to reach criterion and percentage of mice reaching criterion on each day were compared between genotypes.

### Pentylenetetrazol-induced seizures

Behavioural assessment of seizure threshold in mice was performed with injections of 80 mg/kg of pentylenetetrazol (PTZ) as described previously^42,56^. PTZ-induced convulsions were used to gauge susceptibility to primary generalized seizures and as a gross approximation of excitation – inhibitory balance.

### Behavioural Analysis

Data were analyzed in Graphpad PRISM 7.0. Sexes were considered separately with genotype as the fixed factor. All significance levels were set at p < 0.05 and all tests were two-tailed.

## Results

### Model validation

We used RNA sequencing (RNA-seq) on four samples of micro-dissected cerebellum tissue from adult (∼PND145) mice with heterozygous null mutation of *Arid1b* or their WT littermates that had been run through the behavioural experiments. We tested for differential expression across 8,942 genes that were robustly expressed in our samples. We found a total of 1001 differentially expressed (DE) genes in our model with 501 genes meeting an FDR of 0.25 (p < 0.01), 235 genes meeting an FDR of 0.15 (p < 0.004), and 72 genes meeting an FDR of 0.05 (p < 0.0004). Of the total significant DE genes, 607 were downregulated and 394 were upregulated. Almost all DE genes (99.9%) had a subtle change in expression with an absolute log_2_ fold change (FC) of less than 1 (**Figure 1A**). The most significantly DE gene was Somatostatin (log_2_ FC = 0.74, p = 2.92E-09, FDR = 2.61E-05). While not the top gene, we validated that *Arid1b* was differentially expressed in our model (log2 FC = −0.37, p = 0.0037, FDR = 0.14; **Figure 1B**). A heatmap showed clear separation of DE genes between heterozygous *Arid1b* null and WT litter mate samples providing further validation of our genetic model (**Supplementary Figure 1**). Gene set enrichment analysis of Gene Ontology (GO) terms showed enrichment for annotations associated with neuronal morphology and synaptic function in upregulated genes and annotations associated with gene regulation in downregulated genes (**Figure 1C**; **Supplementary Table 1**). We found differential expression of autism-relevant genes in our DE gene list that are also associated with cell adhesion (*Dscam, Reln*) or gene regulation (*Kat2b, Kmt2c, Med13/Med13l, Tbl1xr1, Ubn2*; **Supplementary Table 1**).

**Figure 1.**
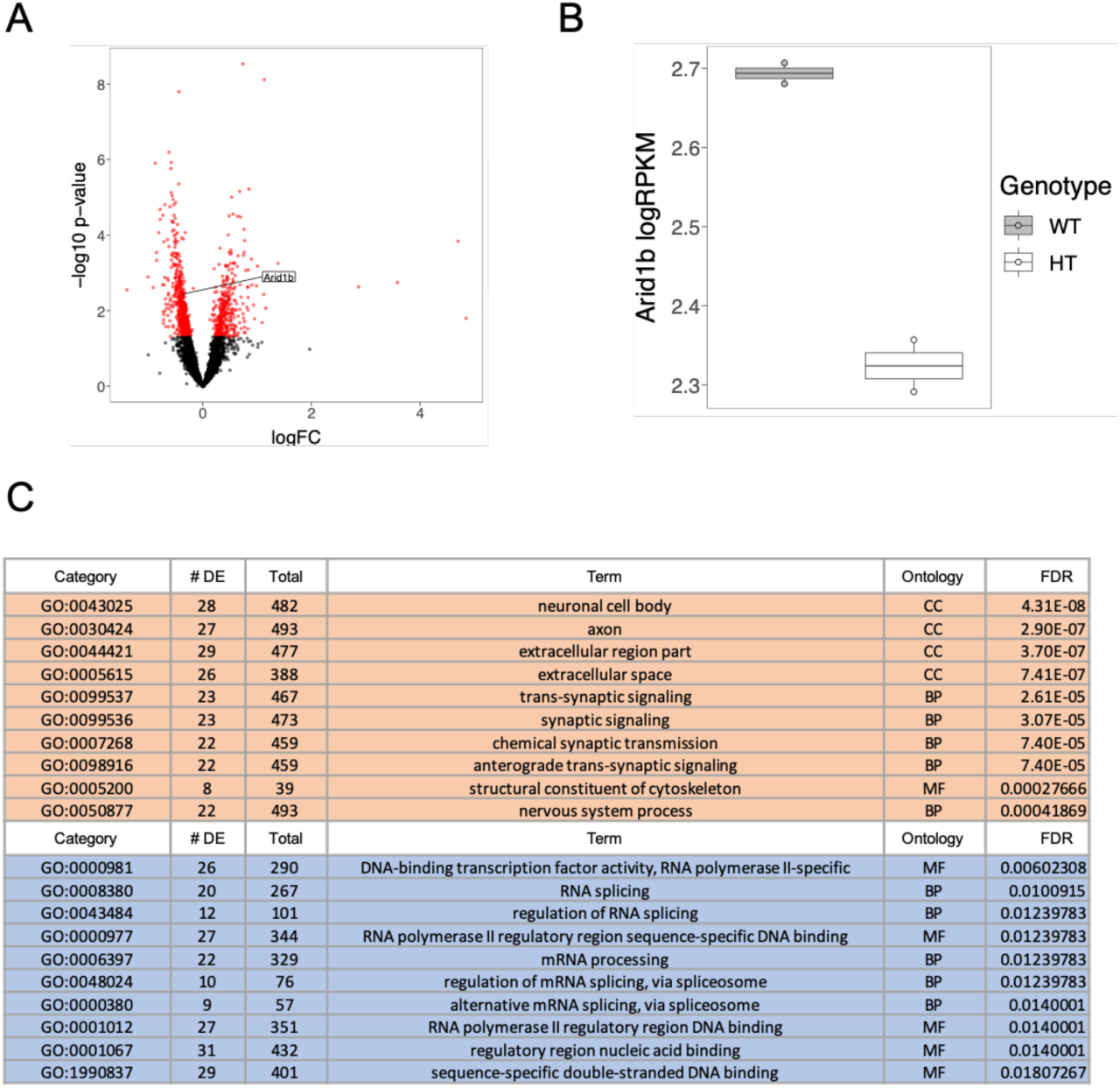
Model Validation. (A) Scatterplot of differentially expressed genes in heterozygous mice. Genes with p-value < 0.05 are in red. (B) Bar plot indicating logRPKM *Arid1b* expression between wild-type (WT) and heterozygous (HT) adult mutants. (C) Table showing select pathways enriched in differentially expressed genes meeting an FDR cutoff of 0.2. Pathways enriched in up-regulated genes are shown in the peach color. Pathways enriched in down-regulated genes are shown in blue. Pathways shown have 500 or fewer genes annotated to their category. Ontologies are biological pathways (BP), molecular function (MF), or cellular component (CC).

### Hydrocephaly

There was an increased incidence of hydrocephaly in the *Arid1b*^*+/-*^ mice compared to other mice in both Sacramento and Toronto colonies. In the time period leading up to this study and others 667 *Arid1b*^*+/-*^ mice were born in the Toronto lab. Of those mice, 21 or 3.1% had to be sacrificed or removed from the analysis due to the hydrocephaly. Those animals were humanely euthanized and not included in our developmental or adult behavioural data. From other colonies in our Toronto based lab, the incidences of hydrocephaly in the mice on C57BL6/N background ranged from 0.2 to 0.3%. In the Sacramento lab, 89 *Arid1b*^*+/-*^ mice were born for this study, 55 of which were used for this study and 7 were excluded or died due to hydrocephalus, a rate of 12.7%, in equal rates between males and females. Of the 82 *Arid1b*^*+/+*^ used in this study, only one was excluded due to hydrocephalus, a rate of 1.2%. Occurrence of hydrocephalus has been previously reported in *Arid1b*^*+/-*^ mice models at rates of 5.5% to 6.6%^17,19^. These rates were mouse models that had severe hydrocephaly and thus were excluded from the study, but it should be noted that the ventricular system as a whole was enlarged in *Arid1b*^*+/-*^ mice throughout development (q=0.005), particularly the lateral ventricles (q=2.0×10^−12^).

### Structural Findings – Adult

At adulthood, the total brain volume of the *Arid1b*^*+/-*^ mutants was observed to be 2.21% smaller than their corresponding WT littermates. However, this did not reach significance (p=0.02, False Discovery Rate (FDR) q=0.37, 420 ± 14 mm^3^ for the *Arid1b*^*+/-*^ mice and 429 ± 12 mm^3^ for the WT, see **Supplementary Table 2**). In the 282 regions examined, 79 of them were significantly different at q<0.05. The main olfactory bulb was significantly smaller (−6.10%, q=0.0007) in size with the olfactory regions within ranging from −5.96 to −10.41% (**Supplementary Table 2**). Similarly, the brainstem was also significantly smaller (−3.13%, q=0.046), due to the smaller hindbrain and pons (−5.53%, q=0.0004 and −4.99%, q=0.0005, respectively). Overall, the white matter throughout the brain was found to be significantly smaller (−3.71%, q=0.046). However, the difference in white matter stemmed from both the cerebellar and cranial nerve fibers, whereas the largest white matter tracts in the mouse brain, the fimbria and corpus callosum, commonly found altered in models of neurodevelopmental disorders, were unaffected (**Supplementary Table 2**). The largest differences, however, were in the cerebellum (−6.77%, q=0.00007), with the cerebellar cortex decreasing by −6.65% (q=0.0001) and the deep cerebellar nuclei (DCN) decreasing by −10.99% (q=4×10^−8^). Relative volume differences were also measured by covarying for total brain volume. The majority of the differences found in relative volume were consistent with absolute volume (**Supplementary Table 2**). However, there were significant increases in the relative volume of the cortex (q=0.01), the striatum (q=0.006), and the corpus callosum in the *Arid1b*^*+/-*^ (q=0.007).

**Figure 2** shows a visual representation of the differences found throughout the brain in the *Arid1b*^*+/-*^ mutants. Sex differences in these results revealed that the *Arid1b*^*+/-*^ mutation appeared to have a stronger effect on the females with 47 of the 282 regions found to be significantly different in the females and only 9 of the 282 regions in the males (**Supplementary Table 2**). However, there was no sex by genotype interaction found in the adult mice indicating that the effect of the mutation was similar in the males and females in adulthood.

**Figure 2.**
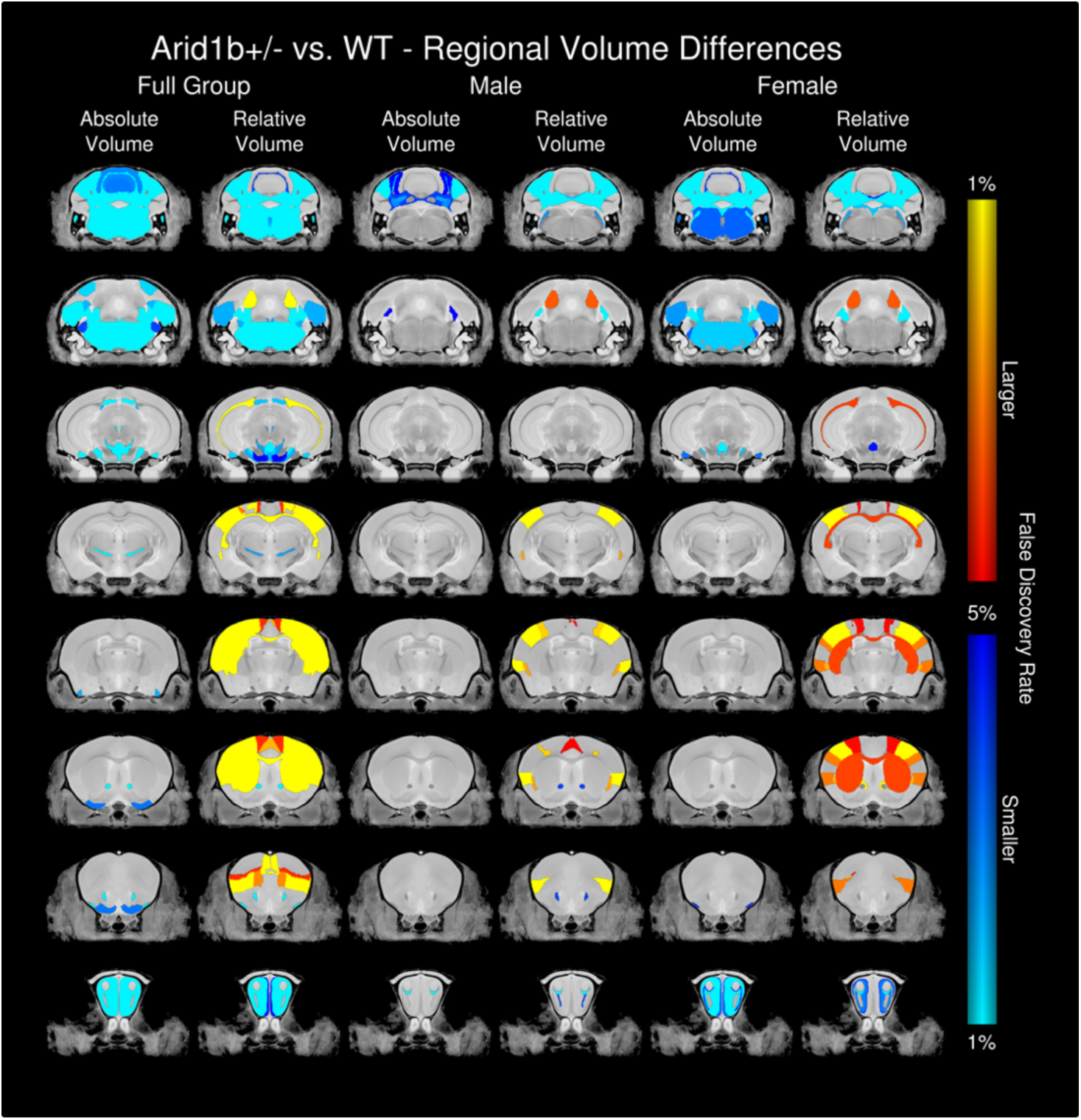
Structural difference in volume found throughout the brain. Shown in both absolute and relative volume in the full group, males, and females. Absolute volume differences are measured in mm^3^, whereas relative volume differences are calculated from a linear model in which the total brain volume is used as a covariate.

### Structural Findings – Longitudinal

The results of the linear mixed effects model showed that *Arid1b*^*+/-*^ (genotype) has a large effect on several structures across the brain (**Figure 3A, Supplementary Table 3**). Across ages there appears to be an overall loss in total brain volume (q=0.02). The strongest differences were seen in the cerebellum and hippocampus. The cerebellum as a whole was found to be smaller in the *Arid1b*^*+/-*^ mice (q=3×10^−8^); this affected all areas of the cerebellum including the vermal regions (q=8×10^−7^), the hemispheric regions (q=0.0004) with strong effects in both the arbor vitae (q=3×10^−11^) and the deep cerebellar nuclei (q=2×10^−9^). In the hippocampal region notable differences were found including an increase in the size of the CA1 (q=0.003), CA2 (q=0.001), CA3 (q=2×10^−8^), and the dentate gyrus (q=6×10^−7^).

**Figure 3.**
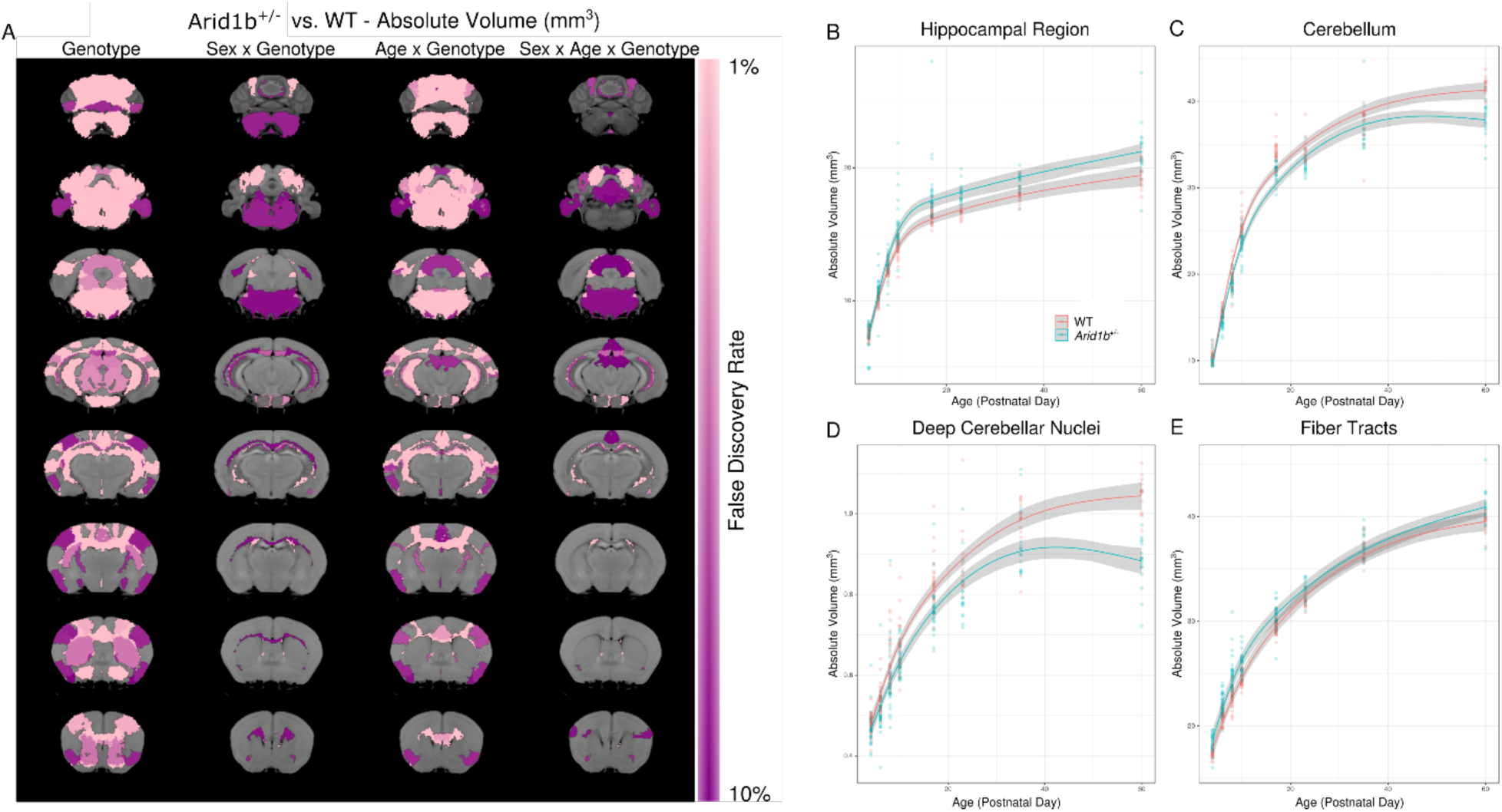
Effects of *Arid1b*^*+/-*^ on the brain throughout development. A) Highlights the results from the linear mixed effects model. The first column represents the main effect of genotype across the study sample. The next highlights the sex by genotype interaction, followed by the age by genotype interaction, and the last column shows the three-way interaction between age, sex, and genotype. The growth rates of the B) hippocampal region, C) cerebellum, D) deep cerebellar nuclei, and E) fiber tracts are shown throughout development. The trend lines shown and the 95% confidence intervals are based on predictions from the linear mixed effects model.

Further investigation revealed a significant interaction with age and genotype, showing that *Arid1b*^*+/-*^ mutants demonstrate different growth rates, mainly highlighting an increased growth rate in the hippocampus (**Figure 3B**, q=0.00008) and a decreased growth rate in the cerebellum (**Figure 3C**, q=0.00004) compared to WT. Differences in growth rate were seen in all areas of the cerebellum including the vermal regions (q=0.0006), hemispheric regions (q=0.03), but also the deep cerebellar nuclei (**Figure 3D**, q=0.0001), and the arbor vitae of the cerebellum (q=4×10^−9^) (**Supplementary Table 3**). In addition to the white matter of the cerebellum there were strong differences in white matter fiber tracts in general (q=0.00009), but outside of the cerebellum white matter the growth rate tended to be increased in the *Arid1b*^*+/-*^ mutants. The growth rates in the cerebellum mirror the overall differences between genotypes in adulthood indicating that all areas of the cerebellum are growing at a slower rate during development resulting in the differences seen in adulthood.

We also observed a significant interaction between sex and genotype, and in turn several differences were also found in the three-way interaction between age, sex, and genotype (**Figure 3A**). The differences between the male and female *Arid1b*^*+/-*^ mutants across development stem from the neuroanatomical differences emerging much earlier in the male than in the female *Arid1b*^*+/-*^ mutants, which happens regardless of whether the differences were larger or smaller in the adult *Arid1b*^*+/-*^ mutants (**Figure 4**). **Figure 4A** highlights the top 50 structures (based on percent difference) for both larger and smaller regions in the *Arid1b*^*+/-*^ mutants versus the WT mice. Both the structures that are smaller (**Figure 4B**) and larger (**Figure 4C**) at PND60 in the *Arid1b*^*+/-*^ mutants diverge earlier in the males, with differences seen as early as PND7. In the females the neuroanatomical differences do not separate from the WT until after PND40 (**Figure 4B and C**).

**Figure 4.**
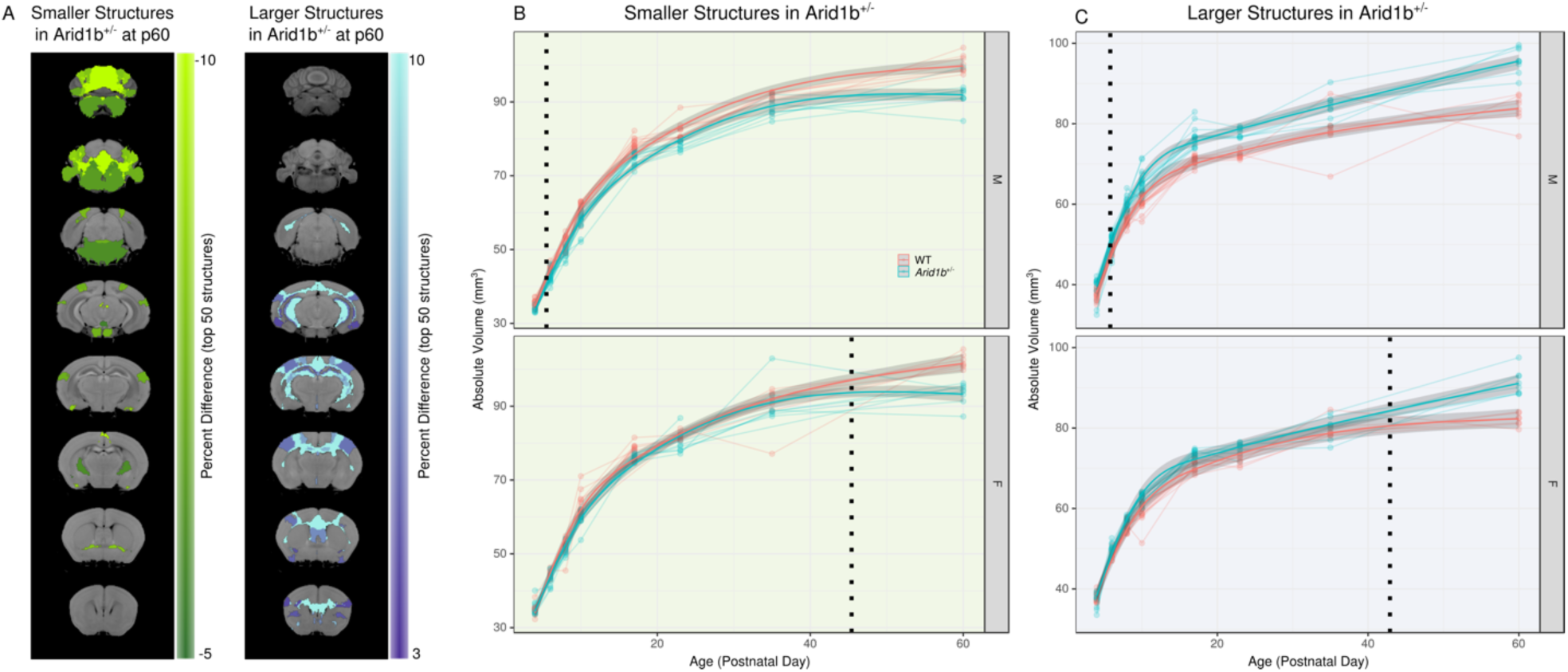
Effects of *Arid1b*^*+/-*^ on the brain throughout development. A) Highlights the top 50 structures on coronal slices that were larger or smaller in the *Arid1b*+/- mouse at PND60. The delayed emergence of the differences in the female *Arid1b*+/- mutants are show for the smaller (B) and larger (C) structures. The dotted line shows when significant differences emerge for the *Arid1b*+/- mice vs. the WT. The trend lines and the 95% confidence intervals shown are based on predictions from the linear mixed effects model.

### Behavioural Findings

*Arid1b*^*+/-*^ pups were impaired on several parameters of developmental milestones including growth, reflexes, and pup ultrasonic vocalization (USVs) emissions, when compared to littermate *Arid1b*^*+/+*^ *controls*. The pups were tested on assays of developmental milestones every other day from postnatal days 2, 4, 6, 8, 10, 12 as previously described^57-59^. USVs were collected by the pup isolation paradigm on days 3, 5, 7, and 9 as previously described^42^. Physical developmental milestones were body weight, body length and head width. Behavioural developmental milestones were righting reflex, negative geotaxis, cliff aversion, circle transverse, forepaw grasping reflex, bar holding, forelimb hang and hindlimb hang. The cut-off latency for timed assays was 30 seconds. **Figure 5A** illustrates isolation-induced USVs over a developmental time course. As expected, a significant effect of age was detected on the rate of calls over time (F_(3, 171)_ = 6.68, p≤0.005, two-way repeated measures ANOVA). Genotype effects across time were also detected (F_(1, 57)_ = 6.67, p≤0.02, one-way repeated measures ANOVA), most prominently at PND3 and 5 (*Arid1b*^*+/-*^ vs. *Arid1b*^*+/-*+^, Bonferroni-Sidak posthoc). Sum of total calls collected across the study were fewer in the *Arid1b*^*+/-*^ (**Figure 5B**; t_(58)_ = 3.08, p≤0.005, student’s t-test), compared to *Arid1b*^*+/+*^ littermates. Core temperature (**Figure 5C**; F_(1, 57)_ = 3.92, NS) was not different by genotype, a variable known to alter pup USVs^60^.

**Figure 5:**
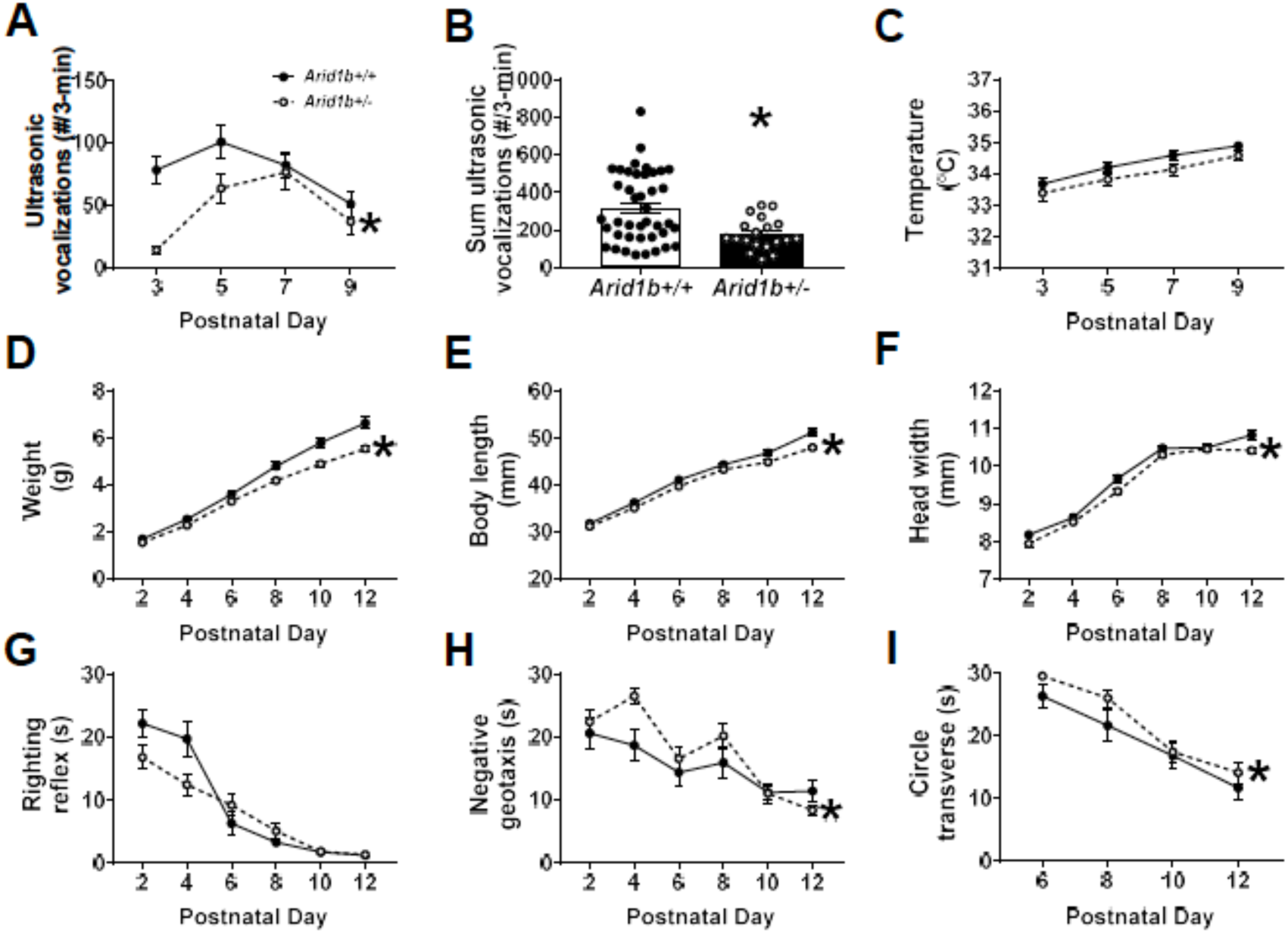
Neonatal ultrasonic vocalization emissions, developmental growth, neurological reflexes and developmental delays in *Arid1b*^*+/-*^. (A) Across neonatal developmental life, *Arid1b*^+/-^ pups displayed abnormal ultrasonic vocalization (USVs) emissions including delay of peak number of calls and decreased total number of USVs on several days compared to *Arid1b*^*+/+*^. (B) When summed, *Arid1b*^+/-^ pups emitted significantly fewer USVs compared to *Arid1b*^*+/+*^ littermate controls. (C) No genotype difference in axillary abdominal temperature was found, confirming fewer USVs were not the result of neonatal hypothermia. (D) *Arid1b*^+/-^ weighed less across development beginning at postnatal 8 and continuously throughout development compared to sex-matched *Arid1b*^*+/+*^. (E) Across neonatal developmental life, *Arid1b*^+/-^ pups were shorter by body length (F) and narrower by head width measurements. (G) While *Arid1b*^+/-^ showed normal latencies to perform the righting reflex, a measure of limb coordination in early life, (H) *Arid1b*^+/-^ had prominent deficits in negative geotaxis (i.e., incline reorientation) and (I) the ability to traverse out of circle by walking. For (A), (B), (C), *Arid1b*^+/+^ N=40, *Arid1b*^+/-^ N=19, for (D-I), *Arid1b*^+/+^ N=18, *Arid1b*^+/-^ N=29. * p<.05 vs *Arid1b*^+/+^ by Repeated Measures Two-way ANOVA or Student’s Unpaired t-test. Error bars represent mean ± SEM.

Body weight, length, and head width were lower, smaller and delayed in *Arid1b*^*+/-*^, respectively (**Figure 5D**; F_(1, 45)_ = 14.8, p≤0.005; **Figure 5E**; F_(1, 45)_ = 14.07, p≤0.005; **Figure 5F**; F_(1, 45)_ = 8.10, p≤0.01, Two-Way Repeated Measures ANOVA), indicating atypical growth and development. Posthoc analysis revealed *Arid1b*^*+/-*^ weighed less on PND6-12, were smaller on PND10-12 and had narrower heads on PND12 vs. *Arid1b*^*+/*+^ (Sidak posthoc for all analysis).

*Arid1b*^*+/-*^ did not demonstrate hypotonia and physical core coordination deficits as they did not have longer latencies to right their bodies dorsally, but longer latencies to reverse from an inclined position in negative geotaxis and time to traverse out of circle were detected in *Arid1b*^*+/-*^, respectively (**Figure 5G**; F_(1, 45)_ = 3.71, p=0.06; **Figure 5H**; F_(1, 45)_ = 4.54, p≤0.05; **Figure 5I**; F_(1, 45)_ = 4.34, p≤0.05, Two-Way Repeated Measures ANOVA), which indicates delayed motoric reflexes, complex limb coordination, and onset of walking skills. Clear divergence in physical metrics were detected in the mid and late ranges of time points collected. As the pups became more fully developed, some metrics were difficult to collect with reliable accuracy at every time point which may contribute to the incomplete penetrance of the phenotypes. However, the strong genotype effect by strong statistic was convincing that *Arid1b*^*+/-*^ diverged by numerous major developmental metrics.

Adult *Arid1b*^*+/-*^ remained smaller and weaker in measurements of size and strength. **Figure 6A** illustrates adult *Arid1*^*+/-*^ weighed less compared to *Arid1b*^*+/+*^ through adulthood during behaviour testing. Body weight was collected across the study and all animals gained mass across all time points PND65, PND100 and PND135 (**Figure 6A**; F_(2, 102)_ = 92.53, p≤0.005, two-way repeated measures ANOVA) but less by the *Arid1b*^*+/-*^ males and females (**Figure 6A**; F_(1, 51)_ = 15.22, p≤0.005, two-way repeated measures ANOVA), compared to *Arid1b*^*+/+*^ sex-matched littermates. *Arid1b*^*+/-*^ showed decreased maximum forelimb force exerted, indicating reduced muscle strength by a forelimb grip strength assay (**Figure 6B**; t_(51)_ = 3.08, p≤0.005, student’s unpaired t-test). Nuanced motoric behaviours, such as gait, balance, posture, stride and motor coordination can be measured using innovative automated treadmill systems, such as DigiGait™ or Catwalk™. Despite their smaller size, *Arid1b*^*+/-*^ did not have genotype differences in stride length (**Figure 6C**; F_(1, 42)_ = 3.09, NS, two-way repeated measures ANOVA) nor stride frequency (**Figure 6D**; F_(1, 43)_ = 2.59, NS, two-way repeated measures ANOVA) in either fore and hind paws. While a decrease in stride length and an increase in stride frequency were expected due to their diminished size, the lack of deficit in these two major parameters indicate motor coordination of the limbs were intact and this phenotype does not confound other complex behaviours.

**Figure 6:**
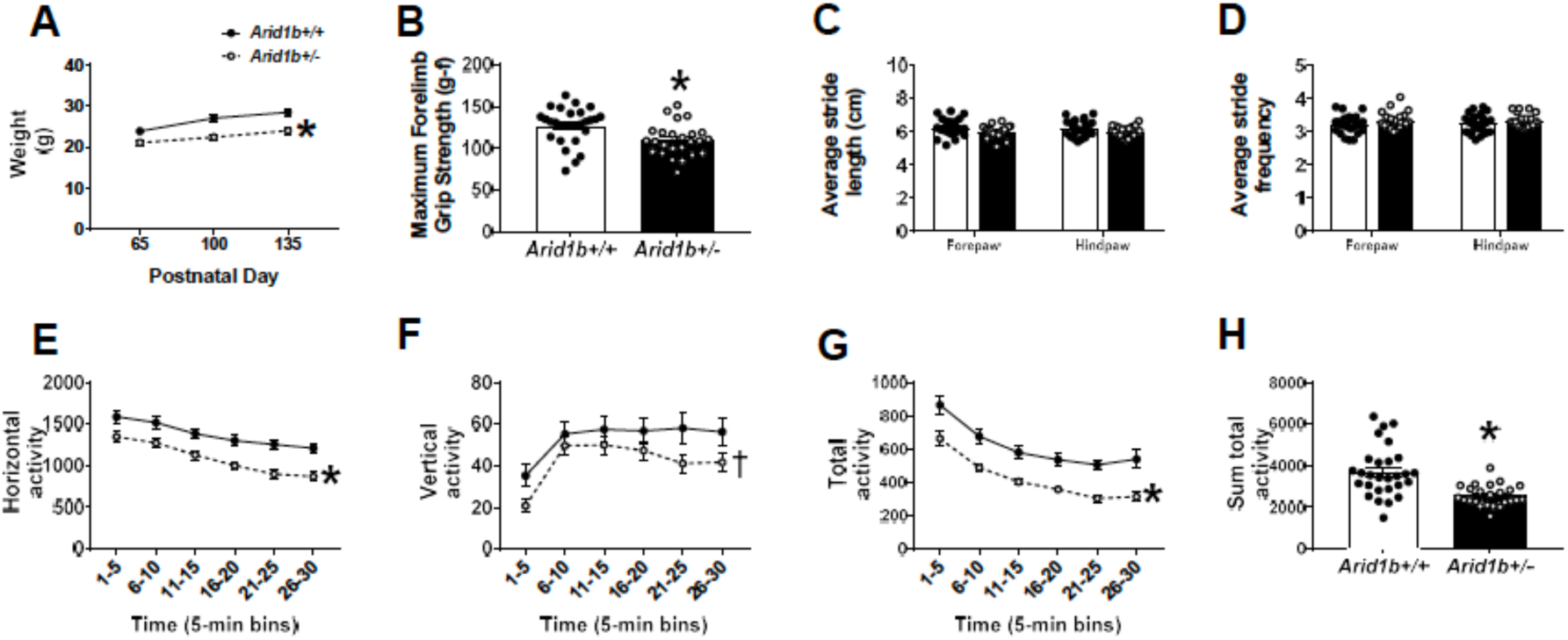
Adult *Arid1b*^*+/*^ have diminished physical size, strength and motor ability. (A) Adult *Arid1*^*+/-*^ weigh less compared to *Arid1b*^*+/+*^ adults and throughout behavioural testing. (B) In a forelimb grip strength assay, *Arid1b*^*+/-*^ showed decreased maximum forelimb force exerted, indicating reduced muscle strength. (C-D) Despite their smaller size, *Arid1b*^*+/-*^ showed normal stride length (C) and stride frequency (D) in fore and hind paws using DigiGait automated gait analysis, indicating normal ambulation and motor coordination of the limbs. (E-H) In open field exploration assay, horizontal, vertical and total activity were recorded for a 30-minute period. Data are shown over time in 5-minute bins. *Arid1b*^*+/-*^ were hypoactive with reduced horizontal activity (E), vertical activity (F) and total activity (G). (H) When summed over the 30-minute session, *Arid1b*^*+/-*^ had less total activity compared to *Arid1b*^*+/+*^ (H), indicating a robust motor deficit. For (A-B), *Arid1b*^*+/+*^ N=28, *Arid1b*^*+/-*^ N=25, for (C-D) *Arid1b*^*+/+*^ N=21, *Arid1b*^*+/+*^ N=19, for (E-H) *Arid1b*^*+/+*^ N=28, *Arid1b*^*+/-*^ N=26. * p<.05 vs *Arid1b*^*+/+*^ by repeated measures two-way ANOVA or student’s unpaired t-test. † p<.09 vs *Arid1b*^*+/+*^ by repeated measures two-way ANOVA. Error bars represent mean ±SEM.

Adult *Arid1b*^*+/-*^ were impaired on multiple parameters of gross motor skills using an open field novel arena assay of locomotion, when compared to littermate *Arid1b*^*+/+*^ sex-matched littermate controls. *Arid1b*^*+/-*^ were hypoactive over a 30-min session. Difference for horizontal and vertical activity and total distance traversed in the novel open field, were significant, representing normal habituation to the novelty of the open field in both genotypes (**Figure 6E-H**; horizontal F_(5, 260)_ = 42.02, p≤0.005; vertical F_(5, 260)_ = 26.06, p≤0.005; total activity F_(5, 260)_ = 52.2, p≤0.005, two-way repeated measures ANOVA). *Arid1b*^*+/-*^ mice were hypoactive by horizontal and vertical activity during all 5 min binned sessions (**Figure 6E**; F_(1, 52)_ = 14.71, p≤0.005; **Figure 6F**; F_(1, 52)_ = 2.80, p≤0.005, two-way repeated measures ANOVA). *Arid1b*^*+/-*^ mice were also hypoactive by total distance traversed over time (**Figure 6G**; F_(1, 52)_ = 20.96, p≤0.005, two-way repeated measures ANOVA) and summed across the 30-minute session (**Figure 6H**; t_(52)_ = 4.58, p≤0.005), indicating a clear, corroborated motor deficit. Behaviours relevant to ASD were assessed using two corroborating assays of social behaviour, as described previously ^10,47-49^. Interestingly, sociability scores from the automated three-chambered social approach task on the chamber time parameter in the *Arid1b*^*+/-*^ and *Arid1b*^*+/+*^ mice were typical and showed significant sociability (**Figure 7A**; two-way repeated measures ANOVA main effect of genotype: F_(1, 51)_ = 3.016, p>0.05; post-hoc novel object vs. novel mouse: *Arid1b*^*+/+*^ p<0.001, *Arid1b*^*+/-*^ p<0.001).

**Figure 7:**
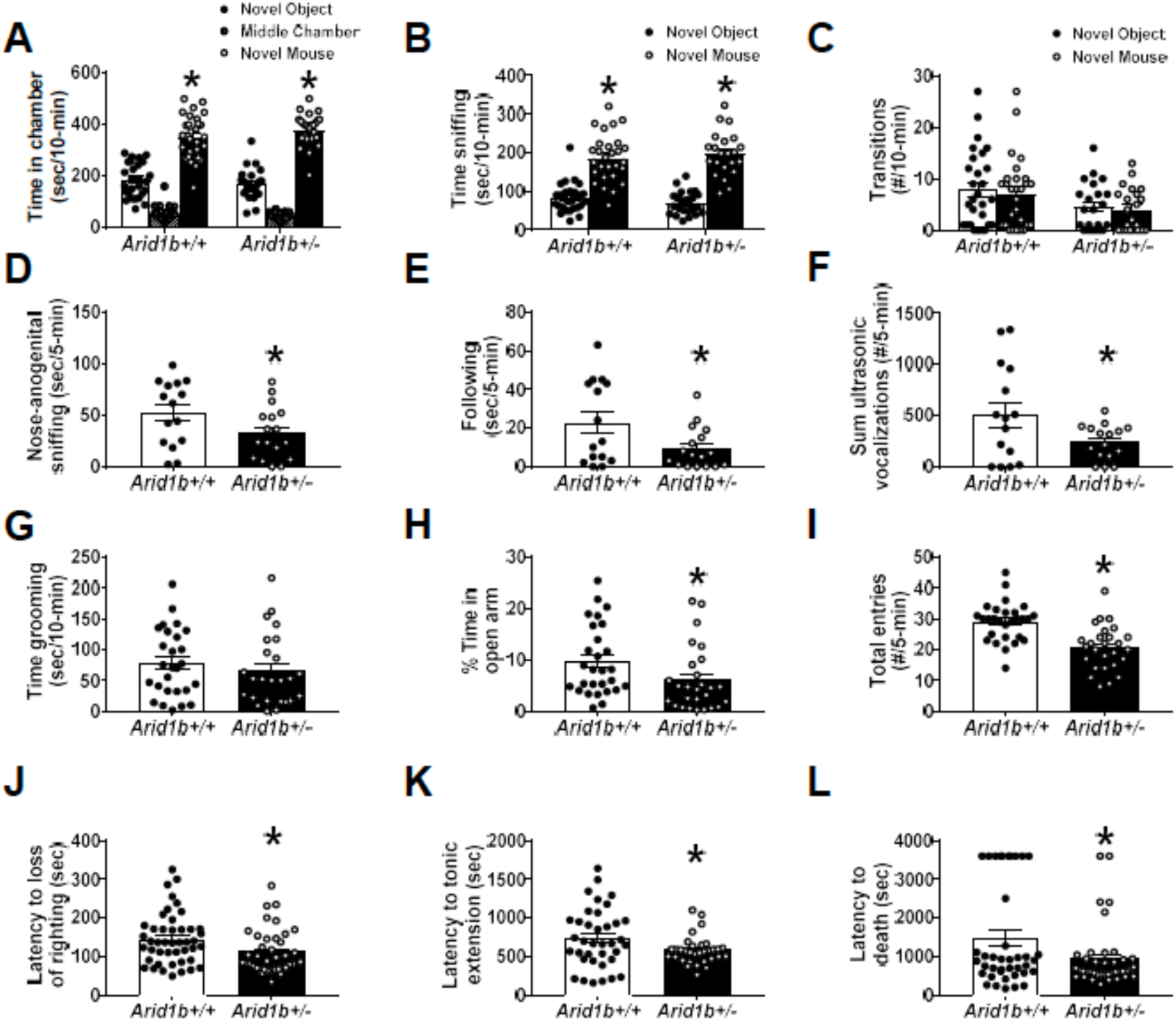
*Arid1b*^+/-^ exhibited mild ASD-relevant social communication phenotypes in the direct reciprocal social interaction and seizure susceptibility but no repetitive behaviour. Three-chambered social approach is a standard measure of social behaviour. It was used as a preliminary evaluation tool for sociability. While both genotypes exhibited normal social approach i.e., they spent more time in the chamber with a novel mouse or time sniffing the novel mouse compared to a novel object), the *Arid1b*^*+/-*^ made fewer total entries during the task. (A) *Arid1b*^*+/-*^ showed no deficits compared to *Arid1b*^*+/+*^ littermates in 3-chambered social approach in time spent in chamber with the novel mouse, (B) time spent sniffing the novel mouse compared to novel object also indicated sociability in both genotypes. (C) *Arid1b*^+/-^ made fewer transitions compared to *Arid1b*^*+/+*^. (D-F) Next, we moved to a more sensitive task to observe social dyad interactions. During male-female social dyad interactions, males are introduced to a WT, novel estrous female for 5 minutes in a novel, clean environment and social investigative and situational behaviours are evaluated and male-emitted ultrasonic vocalizations (USVs) are counted. (D) Adult *Arid1b*^*+/-*^ males spent less time nose-to-anogenital sniffing and (E) less time in a following posture/behaviour, the two main components of this interaction, indicating reduced social behaviour in this task. (F) *Arid1b*^*+/-*^ males also emitted fewer USVs during the dyad interactions compared to *Arid1b*^*+/+*^ male littermates which also suggested impaired social communication. (G) No increased or decreased self-grooming was observed. (H-I) We used a gold standard assay of anxiety, the elevated plus maze, to observe anxiety-like behaviour. (H) *Arid1b*^*+/-*^ spent fewer total seconds on the open arm which usually indicates anxiety-like behaviour. (I) *Arid1b*^*+/*^*-* also showed fewer total entries between arms, which suggests this task may be confounded by the data in Figure 2, the robust motoric deficit. (J-L) With a high concordance of epilepsy in ASD, and excitatory inhibitory balance being a prominent theory, we investigated seizure susceptibility in *Arid1b*^*+/-*^. Mice are injected intraperitoneally with pentylenetetrazol (PTZ) chemoconvulsant, and latency to seizure events including loss of righting, tonic-clonic seizure and death were recorded. (J) *Arid1b*^*+/*-^ showed seizure vulnerability and susceptibility by decreased in time to loss of righting after PTZ administration (K) faster onset to tonic-clonic extension and (L) latency to death, compared to *Arid1b*^*+/+*^. For (A-C), Arid1b^+/+^ N=28, *Arid1b*^*+/-*^ N=25, for (D-F) *Arid1b*^*+/+*^ N=15, *Arid1b*^*+/-*^ N=17, in (G), *Arid1b*^*+/+*^ N=28, *Arid1b*^*+/-*^ N=25, for (G-H), *Arid1b*^*+/+*^ N=28, *Arid1b*^*+/-*^ N=27, for (J-L), *Arid1b*^*+/+*^ N=45, *Arid1b*^*+/-*^ N=40. * p<.05 vs *Arid1b*^*+/+*^ by Repeated Measures Two-Way ANOVA or Student’s Unpaired t-Test. Error bars represent mean ±SEM.

Both *Arid1b*^*+/-*^ and *Arid1b*^*+/+*^ mice exhibited significantly more time social sniffing, which was defined as time spent within 2-cm of the wire cup, with the head facing the wire cup containing the stimulus mouse, as compared to the time spent sniffing the novel object, using the same body point detection settings, (**Figure 7B**; two-way repeated measures ANOVA, main effect of genotype: F_(1, 51)_ = 0.03, p<0.005; *Arid1b*^*+/+*^ p<0.001, *Arid1b*^*+/-*^ p<0.001). *Arid1b*^*+/-*^ mice showed fewer entries into the side chambers (**Figure 7C**; two-way repeated measures ANOVA, main effect of genotype: F_(1, 51)_ = 8.49, p=0.05), indicating that reduced *Arid1b* resulted in less general exploratory activity throughout the 3-chambered apparatus during the social approach assay. However, it did not adversely affect the outcome as both genotypes illustrated intact sociability.

Social deficits were observed on investigative and social parameters in male *Arid1b*^*+/-*^ mice when compared to littermate *Arid1b*^*+/+*^ during the male-female reciprocal dyad social interaction test. **Figure 7D–F** illustrates a detailed examination of male-female social interaction parameters during a session of reciprocal interactions between subject male *Arid1b*^*+/-*^ and *Arid1b*^*+/+*^ mice paired with an unfamiliar estrous B6/NJ female. *Arid1b*^*+/-*^ exhibited less time in nose-to-anogenital sniffing (**Figure 7D**; t_(30)_ = 1.97, p≤0.05, Student’s unpaired t-test) and following behaviour (**Figure 7E**; t_(30)_ = 2.27, p≤0.05, Student’s unpaired t-test). Other parameters collected during the social interaction such as nose to nose sniffing, exploration, and grooming were not significant (nose- to-nose sniffing: p=.55, exploration: p =.06, grooming: p = .59). Once again, as adults, ultrasonic call emissions were fewer in the *Arid1b*^*+/-*^ (**Figure 7F**; t_(29)_ = 2.12, p≤0.005, Student’s unpaired t-test), compared to *Arid1b*^*+/+*^ littermates, which may be attributed to reduced social communication but could also be the result of being smaller, with less pronounced larynx musculature^60-62^.

Self-grooming is one of the most frequently performed behavioural activities in rodents and is a useful measure of repetitive behaviour in models of NDDs with high repetitive behaviours. No difference between subject *Arid1b*^*+/-*^ and *Arid1b*^*+/+*^ mice was observed on repetitive self-grooming behaviour (**Figure 7G**; t_(50)_ = 0.82, NS, Student’s unpaired t-test). No other reported repetitive behaviours, such as circling, jumping or back flipping, were observed during this empty cage observational assay.

Anxiety-like behaviour was assessed in the gold-standard elevated plus-maze. *Arid1b*^*+/-*^ mice spent less time on the arm and had fewer total entries calculated by the open and closed arm entries summed (**Figure 7H**; t_(53)_ = 2.10, p≤0.05 and **Figure 7I**; t_(53)_ = 4.61, p≤0.001, Student’s unpaired t-test), compared to the *Arid1b*^*+/+*^ mice. This observation is consistent with less motoric activity in the *Arid1b*^*+/-*^ mice and confounds a clear interpretation of anxiety-like behaviour in this line, although anxiety metrics indicate this to be a phenotype.

Seizures were provoked using pentylenetetrazol (PTZ; 80 mg/kg, i.p.) in *Arid1b*^*+/-*^ and *Arid1b*^*+/+*^ mice littermates, previously described^56^. Latencies to generalized clonic seizure (loss of righting reflex), tonic extension, and death were collected as a preliminary characterization of seizure, subthreshold epileptiform activity, and imbalances in the excitatory/inhibitory homeostasis. Pentylenetetrazol (PTZ) is a non-competitive GABA_A_ antagonist leading to hyperexcitability^63,64^. *Arid1b*^*+/-*^ mice exhibited faster loss of righting reflexes (**Figure 7J**; t_(83)_ = 2.49, p≤0.02, Student’s unpaired t-test), tonic hindlimb extension (**Figure 7K**; t_(67)_ = 2.11, p≤0.04, Student’s unpaired t-test) and death (**Figure 7L**; t_(77)_ = 2.56, p≤0.02, Student’s unpaired t-test).

Learning and memory abilities were evaluated using tests that are standard and measure simple learning and memory, such as contextual and cued fear conditioning and novel object recognition. Delayed contextual fear conditioning was conducted using automated fear-conditioning chambers as previously described^42^. Fear conditioning to either a cue or a context represents a form of associative learning that has been used in many species. Learning and memory was evaluated using two components, a 24-h contextual component and a 48-h cued fear conditioning. High levels of freezing were observed, subsequent to the conditioned stimulus–unconditioned stimulus pairings, on the training day, across *Arid1b*^*+/+*^ and *Arid1b*^*+/-*^ genotypes (**Figure 8E**; training F_(1, 48)_ = 108.9, p≤0.001; genotype training F_(1, 48)_ = 1.74, NS, Two Way repeated measures ANOVA). No significant effects of genotype on freezing levels were observed 24-hours later during the contextual phase (**Figure 8E**; t_(49)_ = 0.21, NS, Student’s unpaired t-test). High levels of freezing were observed, after the auditory cue stimulus was introduced, on cued conditioning day, across *Arid1b*^*+/+*^ and *Arid1b*^*+/-*^ genotypes indicating no impairments in cued conditioning (**Figure 8F**; effect of cue F_(1, 48)_ = 205.4, p≤0.001; genotype F_(1, 48)_ = 2.86, NS, Two Way repeated measures ANOVA). No significant effects of genotype on freezing levels were observed 24-hours later during the contextual phase (**Figure 8E**; t_(49)_ = 0.21, NS, Student’s unpaired t-test).

**Figure 8:**
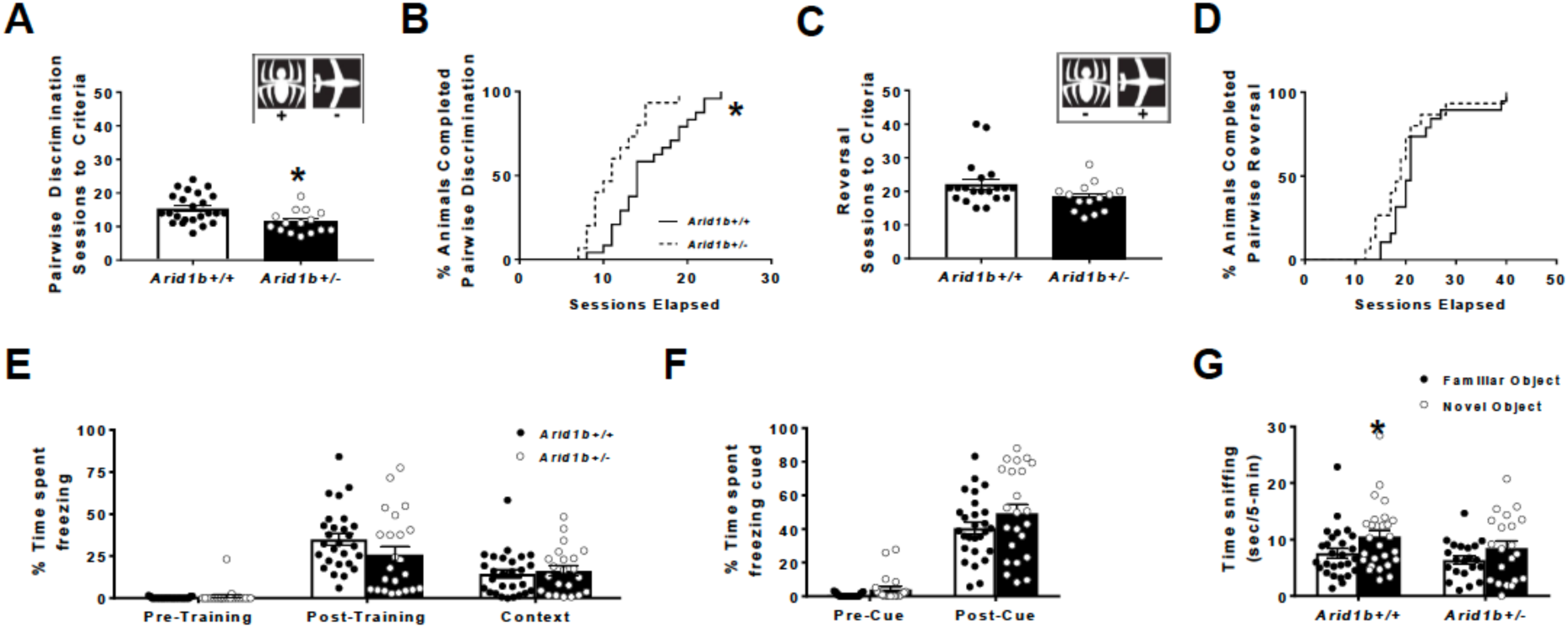
Largely intact learning and memory in *Arid1b*^+/-^. To assess learning and memory and behavioural flexibility in a computerized automated task, that does not rely heavily on motor skills, we utilized a visual pairwise discrimination touchscreen task. Mice are food-restricted and trained to initiate and execute trials by touching their noses to the touchscreen. Mice are then trained to associate an image (a spider or an airplane, A inset) with a food reward. (A) *Arid1b*^*+/-*^ required fewer sessions to reach criteria of 80% accuracy for two consecutive days compared to *Arid1b*^*+/+*^ littermates indicated no deficits in ability to acquire visual discrimination. (B) As a group, *Arid1b*^*+/-*^ completed acquisition of the pairwise discrimination task faster than *Arid1b*^*+/+*^, with larger proportions of mice finishing sooner (survival curve). Once animals achieved task criteria, they underwent reversal of the task where the opposite image now becomes the correct image (C, inset). (C-D) *Arid1b*^*+/-*^ showed no behavioural inflexibility or deficits in learning and memory in pairwise discrimination reversal compared to *Arid1b*^*+/+*^ by (C) no difference in number of sessions required to reach criteria (D) and no difference in proportion of mice reaching criteria of 75% accuracy for two consecutive days (survival curve, D). (E-F) In a canonical learning and memory tasks with less motor involvement, Pavlovian contextual and auditory cue fear conditioning, *Arid1b*^*+/+*^ and Arid1b^+/-^ demonstrated normal associative learning and memory abilities. Learning and memory was assessed by percentage of time spent freezing after associating a context and tone cue with a 0.5 mA foot shock. (E) *Arid1b*^*+/-*^ successfully acquired fear memory during training and showed intact contextual memory of their environment 24 hours later. There was no difference in % freezing between *Arid1b*^*+/+*^ and *Arid1b*^*+/*^. (F) *Arid1b*^*+/+*^ also showed intact cued learning of an auditory tone compared to *Arid1b*^*+/+*^. Recognition memory was tested using a novel object recognition test in an open area. Mice are habituated to an open field arena and 24 hours, familiarized for 10 minutes to two identical objects equally spaced in the arena. After an hour long intertrial interval, one object is replaced with a novel one and mice are allowed to explore for 5 minutes. Time spent sniffing the objects is recorded. Mice exhibit novel object recognition when they spend more time investigating the novel object compared to the familiar object. (G) *Arid1b*^*+/+*^ successfully showed a preference for the novel object and spent more time sniffing the novel object whereas *Arid1b*^*+/-*^ failed to show a preference and spent equal time sniffing both objects. For (A-B), *Arid1b*^*+/+*^ N=24, *Arid1b*^*+/-*^ N=15, for (C-D), *Arid1b*^*+/*+^ N=20, *Arid1b*^*+/-*^ N=14, for (E-F), *Arid1b*^*+/+*^ N=27, *Arid1b*^*+/-*^ N=23, for (G), *Arid1b*^*+/+*^ N=27, *Arid1b*^*+/-*^ N=21. * p<.05 vs *Arid1b*^*+/+*^ by repeated measures Two-Way ANOVA or Student’s Unpaired t-Test. Error bars represent mean ±SEM.

Object recognition is a standard, low stress method performed routinely that can measure short- and long-term memory components. Cognitive abilities as measured by the novel object recognition task were tested in *Arid1b*^*+/+*^ and *Arid1b*^*+/-*^ mice. All automated scores collected via Ethovision were manually confirmed by a trained blinded observer ^40^. *Arid1b*^*+/+*^ spent more time investigating the novel object versus the familiar object, as expected. In contrast, *Arid1b*^*+/+*^ mice did not exhibit typical novel object preference (**Figure 8G**; *Arid1b*^*+/+*^, p < 0.01 and *Arid1b*^*+/-*^ p > 0.05). Both genotypes explored the two identical objects similarly during the automated familiarization phase (data not shown).

We utilized innovative assays of computerized touchscreen assays with high clinical relevance to evaluate visual discrimination and behavioural flexibility attributable to both HPC and cortical circuitry. Preliminary data from our laboratory illustrated the first report utilizing this computer based touchscreen technology in a preclinical model of genetic neurodevelopmental disorders ^51^. Remarkably, **Figure 8A-C** illustrate *Arid1b*^*+/-*^ mice required significantly fewer sessions to a stringent criterion of completing at least 30 trials, at an accuracy of 80% or higher, on two consecutive days, when compared to *Arid1b*^*+/+*^ littermate control subjects. *Arid1b*^*+/-*^ mice needed fewer sessions over more training days to learn to discriminate between two images displayed on the touchscreen (**Figure 8A**; t_(37)_ = 3.07, p≤0.004, Student’s unpaired t-test). Analysis of survival curves, i.e. percentage of mice that reached the 80% accuracy criterion on each training day indicated that the percentage of mice that reached this criterion was significantly higher in *Arid1b*^*+/-*^ than in *Arid1b*^*+/+*^ (**Figure 8B**, X = 8.88, p≤0.003, Gehan Breslow Wilcoxon Chi Square test). These data are in opposition of our hypothesis and illustrate not only that *Arid1b*^*+/-*^ could learn visual discriminations between two juxtaposed computer-generated images, but that they would learn at a faster rate and/or with better accuracy than their *Arid1b*^*+/+*^ littermate control subjects. During reversal, designation of the correct and incorrect stimulus was switched. No genotype effect of *Arid1b*^*+/-*^ was observed in the second, more difficult reversal phase, as shown by sessions to reversal criteria (**Figure 8C**; t_(32)_ = 1.92, NS, Student’s unpaired t-test) and survival curves of mice reaching criterion over time (**Figure 8D**, X = 2.57, NS, Gehan Breslow Wilcoxon Chi Square test).

## Discussion

Deficits in *Arid1b* have been linked to Coffin-Siris syndrome (CSS), autism and intellectual disability. Previous studies have examined various aspects of memory, communication, repetitive behaviours, and social behaviour; however, examination of these behaviours had not been previously assessed throughout development. In this work, haploinsufficiency of *Arid1b*^*+/-*^ resulted in dramatic growth and motor deficits in both the early neonatal stage and adulthood. Similarly, previous studies have only looked at neuroanatomical differences at a single timepoint, missing a possible critical period in development. In this work we performed a whole brain voxelwise analysis across multiple timepoints to assess how the brain changes over time in *Arid1b*^*+/-*^ mice. The original human CSS case studies examined post-mortem brain samples to determine if there were any visible and microstructural differences throughout the brain, and they reported a thinning or absence of the corpus callosum. Therefore, recent mouse studies have only examined one or two regions relevant to the known human neuroanatomical phenotype. Celen et al. reported that the corpus callosum was smaller in adult *Arid1b*^*+/-*^ mice compared to WT^17^, confirming what is seen in a proportion of the human patients. Conversely, while the corpus callosum was not directly assessed in Shibutani et al., they made a general statement that there were no histological abnormalities observed in the *Arid1b*^*+/-*^ compared to WT^19^, but they did not directly assess the corpus callosum volume. In our *ex vivo* whole brain analysis of adult mice we did not find a difference in the size of the corpus callosum, and in fact, we found an increase in the size of the corpus callosum across development in our *in vivo* longitudinal cohort. Additionally, we found an age by genotype interaction affecting the corpus callosum, indicating an increased growth rate in the *Arid1b*^*+/-*^ mutants in not just the corpus callosum but the fiber tracts in general.

The second regional difference reported by Celen et al. was a decrease in the size of the dentate gyrus. In this current assessment, we did not observe any differences in the hippocampal region in either absolute or relative volume in the adult animals, contradicting what was discovered in that study. However, one of the most significant differences in the developmental time course was the hippocampus, and these differences were seen throughout the hippocampus not localized to the dentate gyrus. We observed an increase in the size throughout the *Arid1b*^*+/-*^ hippocampus compared to the WT, which also is contradictory to the previous work. Thus, while *Arid1b*^*+/-*^ does seem to have an effect on the hippocampus, there is inconsistency between studies, which may be attributable to background strain effects (C57BL6/N) compared to the other mouse studies (C57BL6/J).

The above findings are often reported and discussed in recent examinations of CSS patients; however, there is little to no mention of the cerebellar and brain stem findings outside of the original case reports in DeBassio et al.^8^. Interestingly, the cerebellum and brain stem are two of the largest neuroanatomical differences in our study, with the brainstem finding localized to the medulla and pons (**Supplementary Table 2**). These differences confirm in the mouse what was originally reported, where the authors found ectopic collections of neurons in these regions indicating that they were abnormally formed. Moreover, in those original CSS papers, the authors report a Dandy-Walker malformation in several of the children, which is a hypoplasia of the cerebellar vermis with cystic dilation of the fourth ventricle; phenotypes also observed the *Arid1b*^*+/-*^ mouse model used in this work.

Since *ARID1B* has been linked to ASD, the cerebellar differences are not surprising, with cerebellar abnormalities consistently linked to ASD^65,66^. In fact, one of the original MRI findings in human autism patients was an abnormal cerebellar vermis^66^. Further, in mice, the cerebellum has been consistently linked to autism relevant genetic mutations^12,67-69^. In a study which examined 26 different mouse models of autism, differences in the cerebellar volume was one of the most consistent findings across all models^11^. The original CSS findings and the data presented here clearly highlight the cerebellum as an important factor in CSS and ASD in general.

The sex differences in the neuroanatomy throughout development in the *Arid1b*^*+/-*^ mice are quite striking. The delayed onset in females is particularly intriguing, especially since the phenotype becomes similar to the male phenotype in adulthood, indicating different growth rates for each sex. This is consistent with the emergence of sex differences, shown by Qiu et al., in male and female mice in general ^25^. In that study, they show that brain regions that are larger in adult male mice emerge earlier than those that are larger in adult female mice. This is intriguing and indicates a differential growth pattern for male and female mice that is recapitulated here in the *Arid1b*^*+/-*^ mutants, although with the added confound that the differences are caused by a combination of the mutation and sex, not just sex solely. This emphasizes the importance of longitudinal imaging in these autism related mouse populations and could explain some of the differences seen among other preclinical *Arid1b*^*+/-*^ papers. For example, Celen et al. reported neuroanatomical differences were found in 50-day old females, and while the females in our 60-day old adult group do seem to be driving the relative volume difference found in the corpus callosum, it is in the opposite direction. Additionally, the authors of these papers did not specifically look at sex differences in their mice outside of a few measures and either used mixed groups^18^ or only looked at males^19^. The notable sex difference is also intriguing as it shows that the *Arid1b*^*+/-*^ mutation could have a sex dependent effect on the neuroanatomy, which could offer some explanation for the female bias in CSS.

As noted, neuroanatomically, *Arid1b*, appears to have a large effect on the development of the cerebellum regardless of age. The noted decrease in the size of the cerebellum in adulthood, seen in both sexes, appears to develop differentially in the males and females. The mechanism behind this differential development is unknown at this time but does require further investigation in both sexes. Additionally, the hippocampus is affected throughout development, showing both a genotype effect, a sex by genotype interaction, as well as an age by genotype interaction. Interesting in the adult mice, this hippocampal finding is absent, as in there is no noticeable effect of *Arid1b* on the hippocampus in adulthood regardless of sex. This indicates that while the hippocampus has an unusual developmental pattern, it does develop to be the appropriate size by adulthood. The effect of this altered development will need to be investigated further, and it may still lead to differences in adulthood that are undetected at the mesoscopic scale examined here. However, learning and memory behavioural tasks assessed as *Arid1b* mutants did not show any deficits in hippocampal memory tasks as adults, suggesting functional effects, if any, do persist.

Overall, the neuroanatomical phenotype in the *Arid1b*^*+/-*^ mouse appears to be a smaller cerebellum, particularly in the white matter and vermal structures, in conjunction with a larger relative isocortex, striatum, and corpus callosum. This is consistent with several previously published mouse models related to autism, including the *Chd8*^*+/-*10,70,71^, 22q11.2^72^, *Fmr1*^68^, *Itgb3*^73^, *En2*^11^, *Nrxn1a*^11^, and *Shank3*^11^ mouse models. The relationship to the Chd8^+/-^ mouse models is particularly interesting as both mouse models are related to chromatin modification. What this does seem to indicate, however, is that a smaller cerebellum, with a larger cortex and corpus callosum, may delineate a subset of ASD.

As numerous genetic studies implicate *ARID1B* as one of the most frequently mutated NDD genes, it was imperative to functionally characterize our novel *Arid1b*^*+/-*^ mutant mice for translationally relevant phenotypes. Our results report a comprehensive developmental examination of neonatal outcomes including reduced body weight, length, and ultrasonic emissions. One of the earlier *Arid1b*^*+/-*^ models reported by Celen et al., described longer call durations and reduced pitch frequency while our report focused on the standard parameter of call number^17^. While a large number of studies report these vocalizations play a significant role in pup retrieval, and pups’ ability to thrive^74^, there is much debate about their function and their translational relevance in genetic mouse models, made using congenic inbred mice beyond the existence of the calls, especially since in mice these calls are innate over learned^75-79^. The number of call emissions as pups is known to be greater in outbred strains, differ across inbred strains, days of post-development, and cage conditions leaving numerous possibilities for the differing results. Most notably, vocalizations in the Celen et al., study were only collected at postnatal day 4 which does not represent the peak day in pup vocalization production. We chose to investigate vocalizations across a developmental timeline and observed that *Arid1b*^*+/-*^ mice emitted fewer vocalizations over the days recorded and in sum across all days, which indicates a social communication deficit. In addition, we observed a shift in the peak day of vocalization in *Arid1b*^*+/-*^ mice from PND5 in the WT pups to PND7 which may indicate a developmental delay in social communication. While our metrics differed from Celen et al., the gross interpretation is similar between reports in that *Arid1b*^*+/-*^ mutant mice had abnormal pup isolation induced USV^17,71^.

We observed that the reduced body size, length, and head width are evident as early as postnatal day 8, indicating that stunted growth, developmental delay, and failure to thrive seen in the clinical population is robustly recapitulated in the model. Additionally, we expanded developmental motor milestones, similar to translational milestones clinically measured in infants and toddlers, and observed delays and deficits in *Arid1b*^*+/-*^ mutant mice. Interestingly, despite delays in negative geotaxis and traversing out of circle *Arid1b*^*+/-*^ mice had preserved righting reflex. These results could reflect a preservation of muscle tone and limb coordination, but a stunting of motor skill gains due to low muscle strength and smaller size. This hypothesis would be supported by our observations of reduced weight and muscular strength in adults. This dramatic phenotype is especially significant considering the prevalence of short stature, developmental delay, and low motor abilities seen in children with CSS^9^, and will be critical to other NDDs, one example is Prader-Willi Sydrome. These models consistently observe robust and developmental phenotypes that are less rigorous and detectable in adults^40^. It would be interesting to pursue growth factor treatment experiments in the future to determine if these phenotypes are reversible.

Social communication and repetitive behaviour interaction deficits are of particular interest in this model because of its association in large ASD genetic studies. Jung et al., found reduced social interaction in an open field as well as in the sociability and social novelty phases of the standard 3-chamberd approach. Celen et al. found reduced social interaction time in males with a caged juvenile mouse. We observed normal 3-chambered social approach but male *Arid1b*^*+/-*^ mice spent less time engaging in social interactions with novel female mouse in male female social dyad interaction assay, more sensitive than the 3-chambered approach^47,48^. We also corroborated the anxiety-like findings of earlier reports. Other common features of ASD such as repetitive behaviour, measured as self grooming and marble burying (data not shown) did not indicate ASD-relevant phenotypes in our model, differing from the high grooming behaviour in the other models (Celen et al. 2017, Jung et al. 2017). Findings here are mostly similar to the earlier reports, but more subtle in effect size with less findings in the ASD-relevant behaviours. Methodological differences including the test order could account for our inability to see as robust differences as the Jung et al., group. The behavioural battery in that study started with stressful Morris water maze testing and T-maze food restriction while ours followed the standard less stressful to most stressful order, previously published by our group^10,80^. In addition, our sample size was twice the sample size per sex than the Jung study. Another common explanation for the behavioural discrepancies, may be due to differences in background strains used; our model utilizes a C57BL/6N background whereas existing other models were generated on a C57BL/6J background. Our laboratory recently highlighted this subtlety of background strain on seizures, social, and cognitive behaviour^81^.

Deep dives into motor skills beyond control testing in open field have been less reported in recent genetic models of NDD. We often observe, in the NDD literature, broad statements on social and cognitive abilities or deficits in abilities are made when the model mice have substantial reductions in size or basal locomotive activity. It cannot be overstated that these observations, when co-occurring, may be “driven” by size or low motor skills^43,44^. A smaller mouse when tested in large chambers used for social approach, water, and T-mazes will always be at a disadvantage. Here, we had the unique opportunity to observe what impact a roughly 15% size reduction and reduced locomotive activity had on control and sophisticated behavioural outcomes. While *Arid1b*^*+/-*^ are small and weak, even well into adulthood, they have preserved motor coordination, gait abilities, and motor learning abilities as demonstrated by the variety of motor tasks performed (**Figure 4**). *Arid1b*^*+/-*^ showed a deficit in novel object recognition, which aligns with cognitive deficits previously found^18^. However, when tested in tasks that are motor independent, we did not observe this phenotype. This comprehensive look at the full profile of behavioural phenotypes is essential for bridging the gap and improving translational predictability. *Arid1b+/-* mice showed normal contextual and cued memory in a fear conditioning paradigm and in a prefrontal cortex pairwise discrimination task, they showed normal or even slightly better performance compared to WT. By investigating learning and memory abilities that do not rely on high levels of movement and exploration, we avoided potential confounds regarding cognitive deficits. It was unexpected to find that our *Arid1b*^*+/-*^ mice did not show motor-independent cognitive deficits. This was surprising as *ARID1B* is a common gene mutated in patients with intellectual disability and learning difficulties are common in children with CSS and ASD^4,9,82^. This may reflect the limitations with the mouse species as a model system or may reflect the high heterogeneity observed in human population. Previous groups have demonstrated learning and memory deficits albeit the contribution of motor impairment was not corrected or considered in discussion despite its phenotypic presence in both earlier reports^43,83^.

Behavioural seizure susceptibility has not been investigated in the previously published models but considering the relevance and presence of seizures and epilepsy present in both CSS and ASD, we used a proconvulsant to observe seizure susceptibility^84^. We observed a strong seizure susceptibility phenotype in *Arid1b*^*+/-*^ mice by numerous metrics including reduced latency to loss of righting, tonic hindlimb extension, and faster death. As are *Arid1b*^*+/-*^ mice were functionally susceptible to PTZ-induced seizures, related to GABAergic neuron plasticity, we attribute this to an excitatory-inhibition imbalance, as other new genetic models of NDD have also exhibited^85-87^. This hypothesis corroborates histological analysis done by Jung et al.^18^.

Coffin-Siris Syndrome is more frequently identified in females than in males; therefore, we investigated sex differences in all behavioural tasks. There was no difference in the number of heterozygous females born compared to males. There were additionally no significant differences in any of the adult behavioural tasks which aligns with adult imaging results and the two other published models^17,18^. However, it is intriguing that we did not identify any sex differences in developmental trajectory tasks different from the clear divergence in neural patterns observed using MR imaging.

Developmental delay appears to be one of the hallmark characteristics found in the *Arid1b*^*+/-*^ mice, which is not just seen with an overall total brain volume loss throughout development, but also with the reduced body weight, length, and head width seen as early as PND8. Specific areas of the brain, however, are more delayed than others. The cerebellum, which is significantly smaller in adulthood, also displays a slower growth rate compared to WT. This reduced cerebellar volume as well as differences seen across development in the primary motor cortex (**Supplementary Table 2**) may be responsible for the motor deficits seen in the *Arid1b*^*+/-*^ mice including their overall decreased activity (**Figure 6H**). The hippocampus on the other hand was found to have an increase growth rate over time, which could be related to the mostly intact learning and memory, and in some cases improvement, seen in motor-independent cognitive tasks. One of the regions closely associated with repetitive behaviours tends to be the striatum^88^, particularly the caudoputamen, in this study we do not see any differences in the adult *ex vivo* group in this key region but do see an increase in the size of the caudoputamen throughout development. As the *Arid1b*^*+/-*^ mice do not show ASD associated repetitive behaviours it is possible that this increase in size of the caudoputamen is preservative for the *Arid1b*^*+/-*^ mice. These statements are speculative at this point, however, as the behaviour and neuroanatomical measures were done on different cohorts of animals; therefore, future examination of these comparisons is warranted.

## Summary

It is clear that the *Arid1b*^*+/-*^ mutation has an effect on both the brain and the behaviour in mice. We have shown dramatic growth and motor deficits throughout development and have confirmed several previously established behavioural phenotypes in the mouse, indicating that the *Arid1b*^*+/-*^ mouse is a good model for neurodevelopmental disorders such as ASD and CSS. While we have been unable to confirm the corpus callosum phenotype that has been previously reported in the mouse, and, in fact, show the opposite effect throughout development, we have highlighted a novel cerebellar phenotype, consistent with that observed in human patients. In addition, the data shows the benefits of examining both sexes throughout development, as several differences reported here would not have been discovered in a single sex, single timepoint study. The delayed emergence of the neuroanatomical phenotype in the female *Arid1b*^*+/-*^ mutants is worth further investigation, especially since it mirrors the emergence of neuroanatomical sex differences. The implications of this delayed onset in females with neurodevelopmental disorders could be far reaching.

## References

1 Banerjee-Basu S, Packer A. SFARI Gene: an evolving database for the autism research community. Dis Model Mech. 2010; 3: 133–135.

2 Ronan JL, Wu W, Crabtree GR. From neural development to cognition: unexpected roles for chromatin. Nature Reviews Genetics 2015 17:1 2013; 14: 347–359.

3 Nord AS, Roeb W, Dickel DE, Walsh T, Kusenda M, O’Connor KL et al. Reduced transcript expression of genes affected by inherited and de novo CNVs in autism. Eur J Hum Genet 2011; 19: 727–731.

4 Halgren C, Kjaergaard S, Bak M, Hansen C, El-Schich Z, Anderson CM et al. Corpus callosum abnormalities, intellectual disability, speech impairment, and autism in patients with haploinsufficiency of ARID1B. Clin Genet 2012; 82: 248–255.

5 Coffin GS, Siris E. Mental retardation with absent fifth fingernail and terminal phalanx. Am J Dis Child 1970; 119: 433–439.

6 Tunnessen WW, McMillan JA, Levin MB. The Coffin-Siris syndrome. Am J Dis Child 1978; 132: 393–395.

7 Imai T, Hattori H, Miyazaki M, Higuchi Y, Adachi S, Nakahata T. Dandy-Walker variant in Coffin-Siris syndrome. Am J Med Genet 2001; 100: 152–155.

8 DeBassio WA, Kemper TL, Knoefel JE. Coffin-Siris syndrome. Neuropathologic findings. Arch Neurol 1985; 42: 350–353.

9 van der Sluijs PJ, Jansen S, Vergano SA, Adachi-Fukuda M, Alanay Y, AlKindy A et al. The ARID1B spectrum in 143 patients: from nonsyndromic intellectual disability to Coffin-Siris syndrome. Genet Med 2018; : 1.

10 Gompers AL, Su-Feher L, Ellegood J, Copping NA, Riyadh MA, Stradleigh TW et al. Germline Chd8 haploinsufficiency alters brain development in mouse. Nat Neurosci 2017; 20: 1062–1073.

11 Ellegood J, Anagnostou E, Babineau BA, Crawley JN, Lin L, Genestine M et al. Clustering autism: using neuroanatomical differences in 26 mouse models to gain insight into the heterogeneity. Mol Psychiatry 2015; 20: 118–125.

12 Ellegood J, Lerch JP, Henkelman RM. Brain abnormalities in a Neuroligin3 R451C knockin mouse model associated with autism. Autism Res 2011; 4: 368–376.

13 Zerbi V, Ielacqua GD, Markicevic M, Haberl MG, Ellisman MH, A-Bhaskaran A et al. Dysfunctional Autism Risk Genes Cause Circuit-Specific Connectivity Deficits With Distinct Developmental Trajectories. Cereb Cortex 2018; 28: 2495–2506.

14 Horder J, Petrinovic MM, Mendez MA, Bruns A, Takumi T, Spooren W et al. Glutamate and GABA in autism spectrum disorder-a translational magnetic resonance spectroscopy study in man and rodent models. Transl Psychiatry 2018; 8: 106.

15 Bertram J, Koschützke L, Pfannmöller JP, Esche J, van Diepen L, Kuss AW et al. Morphological and behavioral characterization of adult mice deficient for SrGAP3. Cell Tissue Res 2016; 366: 1–11.

16 Ellegood J, Crawley JN. Behavioral and Neuroanatomical Phenotypes in Mouse Models of Autism. Neurotherapeutics 2015; 12: 521–533.

17 Celen C, Chuang J-C, Luo X, Nijem N, Walker AK, Chen F et al. Arid1b haploinsufficient mice reveal neuropsychiatric phenotypes and reversible causes of growth impairment. Elife 2017; 6: 2081.

18 Jung E-M, Moffat JJ, Liu J, Dravid SM, Gurumurthy CB, Kim W-Y. Arid1b haploinsufficiency disrupts cortical interneuron development and mouse behavior. Nat Neurosci 2017; 20: 1694–1707.

19 Shibutani M, Horii T, Shoji H, Morita S, Kimura M, Terawaki N et al. Arid1b Haploinsufficiency Causes Abnormal Brain Gene Expression and Autism-Related Behaviors in Mice. Int J Mol Sci 2017; 18: 1872.

20 Cahill LS, Laliberté CL, Ellegood J, Spring S, Gleave JA, Eede MCV et al. Preparation of fixed mouse brains for MRI. Neuroimage 2012; 60: 933–939.

21 Lerch JP, Sled JG, Henkelman RM. MRI phenotyping of genetically altered mice. Methods Mol Biol 2011; 711: 349–361.

22 de Guzman AE, Wong MD, Gleave JA, Nieman BJ. Variations in post-perfusion immersion fixation and storage alter MRI measurements of mouse brain morphometry. Neuroimage 2016; 142: 687–695.

23 Bock NA, Konyer NB, Henkelman RM. Multiple-mouse MRI. Magn Reson Med 2003; 49: 158–167.

24 Spencer Noakes TL, Henkelman RM, Nieman BJ. Partitioning k-space for cylindrical three-dimensional rapid acquisition with relaxation enhancement imaging in the mouse brain. NMR Biomed 2017; 30. doi:10.1002/nbm.3802.

25 Qiu LR, Fernandes DJ, Szulc-Lerch KU, Dazai J, Nieman BJ, Turnbull DH et al. Mouse MRI shows brain areas relatively larger in males emerge before those larger in females. Nat Commun 2018; 9: 2615.

26 Szulc KU, Lerch JP, Nieman BJ, Bartelle BB, Friedel M, Suero-Abreu GA et al. 4D MEMRI atlas of neonatal FVB/N mouse brain development. Neuroimage 2015; 118: 49–62.

27 Nieman BJ, Szulc KU, Turnbull DH. Three-dimensional, in vivo MRI with self-gating and image coregistration in the mouse. Magn Reson Med 2009; 61: 1148–1157.

28 Collins DL, Neelin P, Peters TM, Evans AC. Automatic 3D intersubject registration of MR volumetric data in standardized Talairach space. J Comput Assist Tomogr 1994; 18: 192–205.

29 Avants BB, Epstein CL, Grossman M, Gee JC. Symmetric diffeomorphic image registration with cross-correlation: evaluating automated labeling of elderly and neurodegenerative brain. Med Image Anal 2008; 12: 26–41.

30 Avants BB, Tustison NJ, Song G, Cook PA, Klein A, Gee JC. A reproducible evaluation of ANTs similarity metric performance in brain image registration. Neuroimage 2011; 54: 2033–2044.

31 Lerch JP, Carroll JB, Spring S, Bertram LN, Schwab C, Hayden MR et al. Automated deformation analysis in the YAC128 Huntington disease mouse model. Neuroimage 2008; 39: 32–39.

32 Nieman BJ, Flenniken AM, Adamson SL, Henkelman RM, Sled JG. Anatomical phenotyping in the brain and skull of a mutant mouse by magnetic resonance imaging and computed tomography. Physiol Genomics 2006; 24: 154–162.

33 Dorr AE, Lerch JP, Spring S, Kabani N, Henkelman RM. High resolution three-dimensional brain atlas using an average magnetic resonance image of 40 adult C57Bl/6J mice. Neuroimage 2008; 42: 60–69.

34 Steadman PE, Ellegood J, Szulc KU, Turnbull DH, Joyner AL, Henkelman RM et al. Genetic effects on cerebellar structure across mouse models of autism using a magnetic resonance imaging atlas. Autism Res 2014; 7: 124–137.

35 Ullmann JFP, Watson C, Janke AL, Kurniawan ND, Reutens DC. A segmentation protocol and MRI atlas of the C57BL/6J mouse neocortex. Neuroimage 2013; 78: 196–203.

36 Richards K, Watson C, Buckley RF, Kurniawan ND, Yang Z, Keller MD et al. Segmentation of the mouse hippocampal formation in magnetic resonance images. Neuroimage 2011; 58: 732–740.

37 Genovese CR, Lazar NA, Nichols T. Thresholding of statistical maps in functional neuroimaging using the false discovery rate. Neuroimage 2002; 15: 870–878.

38 Satterthwaite FE. Synthesis of variance. Psychometrika 1941; 6: 309–316.

39 Fox WM. Reflex-ontogeny and behavioural development of the mouse. Anim Behav 1965; 13: 234–241.

40 Adhikari A, Copping NA, Onaga B, Pride MC, Coulson RL, Yang M et al. Cognitive deficits in the Snord116 deletion mouse model for Prader-Willi syndrome. Neurobiol Learn Mem 2018. doi:10.1016/j.nlm.2018.05.011.

41 Flannery BM, Silverman JL, Bruun DA, Puhger KR, McCoy MR, Hammock BD et al. Behavioral assessment of NIH Swiss mice acutely intoxicated with tetramethylenedisulfotetramine. Neurotoxicol Teratol 2015; 47: 36–45.

42 Copping NA, Christian SGB, Ritter DJ, Islam MS, Buscher N, Zolkowska D et al. Neuronal overexpression of Ube3a isoform 2 causes behavioral impairments and neuroanatomical pathology relevant to 15q11.2-q13.3 duplication syndrome. Hum Mol Genet 2017; 26: 3995–4010.

43 Sukoff Rizzo SJ, Silverman JL. Methodological Considerations for Optimizing and Validating Behavioral Assays. Curr Protoc Mouse Biol 2016; 6: 364–379.

44 Silverman JL, Ellegood J. Behavioral and neuroanatomical approaches in models of neurodevelopmental disorders: opportunities for translation. Curr Opin Neurol 2018; 31: 126–133.

45 Sashindranath M, Daglas M, Medcalf RL. Evaluation of gait impairment in mice subjected to craniotomy and traumatic brain injury. Behavioural Brain Research 2015; 286: 33–38.

46 Gulinello M, Mitchell HA, Chang Q, Timothy O’Brien W, Zhou Z, Abel T et al. Rigor and reproducibility in rodent behavioral research. Neurobiol Learn Mem 2018. doi:10.1016/j.nlm.2018.01.001.

47 Silverman JL, Pride MC, Hayes JE, Puhger KR, Butler-Struben HM, Baker S et al. GABAB Receptor Agonist R-Baclofen Reverses Social Deficits and Reduces Repetitive Behavior in Two Mouse Models of Autism. Neuropsychopharmacology 2015; 40: 2228–2239.

48 Dhamne SC, Silverman JL, Super CE, Lammers SHT, Hameed MQ, Modi ME et al. Replicable in vivo physiological and behavioral phenotypes of the Shank3B null mutant mouse model of autism. Mol Autism 2017; 8: 26.

49 Yang M, Silverman JL, Crawley JN. Automated three-chambered social approach task for mice. Curr Protoc Neurosci 2011; Chapter 8: Unit 8.26.

50 Bales KL, Solomon M, Jacob S, Crawley JN, Silverman JL, Larke RH et al. Long-term exposure to intranasal oxytocin in a mouse autism model. Transl Psychiatry 2014; 4: e480–e480.

51 Copping NA, Berg EL, Foley GM, Schaffler MD, Onaga BL, Buscher N et al. Touchscreen learning deficits and normal social approach behavior in the Shank3B model of Phelan-McDermid Syndrome and autism. Neuroscience 2017; 345: 155–165.

52 Scattoni ML, Ricceri L, Crawley JN. Unusual repertoire of vocalizations in adult BTBR T+tf/J mice during three types of social encounters. Genes, Brain and Behavior 2011; 10: 44–56.

53 Brigman JL, Powell EM, Mittleman G, Young JW. Examining the genetic and neural components of cognitive flexibility using mice. Physiol Behav 2012; 107: 666–669.

54 Brigman JL, Daut RA, Wright T, Gunduz-Cinar O, Graybeal C, Davis MI et al. GluN2B in corticostriatal circuits governs choice learning and choice shifting. Nat Neurosci 2013; 16: 1101–1110.

55 Horner AE, Heath CJ, Hvoslef-Eide M, Kent BA, Kim CH, Nilsson SRO et al. The touchscreen operant platform for testing learning and memory in rats and mice. Nat Protoc 2013; 8: 1961–1984.

56 Dhir A. Pentylenetetrazol (PTZ) Kindling Model of Epilepsy. John Wiley & Sons, Inc.: Hoboken, NJ, USA, 2001 doi:10.1002/0471142301.ns0937s58.

57 Scattoni ML, Gandhy SU, Ricceri L, Crawley JN. Unusual repertoire of vocalizations in the BTBR T+tf/J mouse model of autism. PLoS ONE 2008; 3: e3067.

58 Yang M, Abrams DN, Zhang JY, Weber MD, Katz AM, Clarke AM et al. Low sociability in BTBR T+tf/J mice is independent of partner strain. Physiol Behav 2012. doi:10.1016/j.physbeh.2011.12.025.

59 Wöhr M, Silverman JL, Scattoni ML, Turner SM, Harris MJ, Saxena R et al. Developmental delays and reduced pup ultrasonic vocalizations but normal sociability in mice lacking the postsynaptic cell adhesion protein neuroligin2. Behavioural Brain Research 2013; 251: 50–64.

60 Branchi I, Santucci D, Alleva E. Analysis of ultrasonic vocalizations emitted by infant rodents. Curr Protoc Toxicol 2006; Chapter 13: Unit13.12.

61 Branchi I, Santucci D, Alleva E. Ultrasonic vocalisation emitted by infant rodents: a tool for assessment of neurobehavioural development. Behavioural Brain Research 2001; 125: 49–56.

62 Branchi I, Santucci D, Vitale A, Alleva E. Ultrasonic vocalizations by infant laboratory mice: a preliminary spectrographic characterization under different conditions. Dev Psychobiol 1998; 33: 249–256.

63 Rocha L, Briones M, Ackermann RF, Anton B, Maidment NT, Evans CJ et al. Pentylenetetrazol-induced kindling: early involvement of excitatory and inhibitory systems. Epilepsy Res 1996; 26: 105–113.

64 Macdonald RL, Barker JL. Pentylenetetrazol and penicillin are selective antagonists of GABA-mediated post-synaptic inhibition in cultured mammalian neurones. Nature 1977; 267: 720–721.

65 Saitoh O, Courchesne E. Magnetic resonance imaging study of the brain in autism. Psychiatry Clin Neurosci 1998; 52 Suppl: S219–22.

66 Courchesne E, Yeung-Courchesne R, Press GA, Hesselink JR, Jernigan TL. Hypoplasia of cerebellar vermal lobules VI and VII in autism. N Engl J Med 1988; 318: 1349–1354.

67 Allemang-Grand R, Ellegood J, Spencer Noakes L, Ruston J, Justice M, Nieman BJ et al. Neuroanatomy in mouse models of Rett syndrome is related to the severity of Mecp2 mutation and behavioral phenotypes. Mol Autism 2017; 8: 32.

68 Ellegood J, Pacey LK, Hampson DR, Lerch JP, Henkelman RM. Anatomical phenotyping in a mouse model of fragile X syndrome with magnetic resonance imaging. Neuroimage 2010; 53: 1023–1029.

69 Stoodley CJ, D’Mello AM, Ellegood J, Jakkamsetti V, Liu P, Nebel MB et al. Altered cerebellar connectivity in autism and cerebellar-mediated rescue of autism-related behaviors in mice. Nat Neurosci 2017; 20: 1744–1751.

70 Suetterlin P, Hurley S, Mohan C, Riegman KLH, Pagani M, Caruso A et al. Altered Neocortical Gene Expression, Brain Overgrowth and Functional Over-Connectivity in Chd8 Haploinsufficient Mice. Cereb Cortex 2018; 28: 2192–2206.

71 Jung H, Park H, Choi Y, Kang H, Lee E, Kweon H et al. Sexually dimorphic behavior, neuronal activity, and gene expression in Chd8-mutant mice. Nat Neurosci 2018; 21: 1218–1228.

72 Ellegood J, Markx S, Lerch JP, Steadman PE, Genç C, Provenzano F et al. Neuroanatomical phenotypes in a mouse model of the 22q11.2 microdeletion. Mol Psychiatry 2014; 19: 99–107.

73 Ellegood J, Henkelman RM, Lerch JP. Neuroanatomical Assessment of the Integrin β3 Mouse Model Related to Autism and the Serotonin System Using High Resolution MRI. Front Psychiatry 2012; 3: 37.

74 Scattoni ML. Special interest section on mouse ultrasonic vocalizations. Genes, Brain and Behavior 2011; 10: 1–3.

75 Portfors CV, Perkel DJ. The role of ultrasonic vocalizations in mouse communication. Curr Opin Neurobiol 2014; 28: 115–120.

76 Kikusui T, Nakanishi K, Nakagawa R, Nagasawa M, Mogi K, Okanoya K. Cross fostering experiments suggest that mice songs are innate. PLoS ONE 2011; 6: e17721.

77 Kalcounis-Rueppell MC, Petric R, Briggs JR, Carney C, Marshall MM, Willse JT et al. Differences in ultrasonic vocalizations between wild and laboratory California mice (Peromyscus californicus). PLoS ONE 2010; 5: e9705.

78 Wang H, Liang S, Burgdorf J, Wess J, Yeomans J. Ultrasonic vocalizations induced by sex and amphetamine in M2, M4, M5 muscarinic and D2 dopamine receptor knockout mice. PLoS ONE 2008; 3: e1893.

79 Ferhat A-T, Le Sourd A-M, de Chaumont F, Olivo-Marin J-C, Bourgeron T, Ey E. Social communication in mice--are there optimal cage conditions? PLoS ONE 2015; 10: e0121802.

80 Silverman JL, Yang M, Lord C, Crawley JN. Behavioural phenotyping assays for mouse models of autism. Nat Rev Neurosci 2010; 11: 490–502.

81 Copping NA, Adhikari A, Petkova SP, Silverman JL. Genetic backgrounds have unique seizure response profiles and behavioral outcomes following convulsant administration. Epilepsy Behav 2019; 101: 106547.

82 Hoyer J, Ekici AB, Endele S, Popp B, Zweier C, Wiesener A et al. Haploinsufficiency of ARID1B, a member of the SWI/SNF-a chromatin-remodeling complex, is a frequent cause of intellectual disability. Am J Hum Genet 2012; 90: 565–572.

83 Silverman JL, Nithianantharajah J, Der-Avakian A, Young JW, Sukoff Rizzo SJ. Lost in Translation: At the crossroads of face validity and translational utility of behavioural assays in animal models for the development of therapeutics. Neurosci Biobehav Rev. (in press)

84 Adam MP, Ardinger HH, Pagon RA, Wallace SE, Bean LJ, Stephens K et al. Coffin-Siris Syndrome. 1993.

85 Souchet B, Guedj F, Sahún I, Duchon A, Daubigney F, Badel A et al. Excitation/inhibition balance and learning are modified by Dyrk1a gene dosage. Neurobiology of Disease 2014; 69: 65–75.

86 Golden CE, Buxbaum JD, De Rubeis S. Disrupted circuits in mouse models of autism spectrum disorder and intellectual disability. Curr Opin Neurobiol 2018; 48: 106–112.

87 Antoine MW, Langberg T, Schnepel P, Feldman DE. Increased Excitation-Inhibition Ratio Stabilizes Synapse and Circuit Excitability in Four Autism Mouse Models. Neuron 2019; 101: 648–661.e4.

88 Ellegood J, Babineau BA, Henkelman RM, Lerch JP, Crawley JN. Neuroanatomical analysis of the BTBR mouse model of autism using magnetic resonance imaging and diffusion tensor imaging. Neuroimage 2013; 70: 288–300.

