## Supplementary figures and images for "Neuroanatomy and Behaviour in Mice with a Haploinsufficiency of AT-Rich Interactive Domain 1B (ARID1B) Throughout Development"

### Supplementary Figure 1

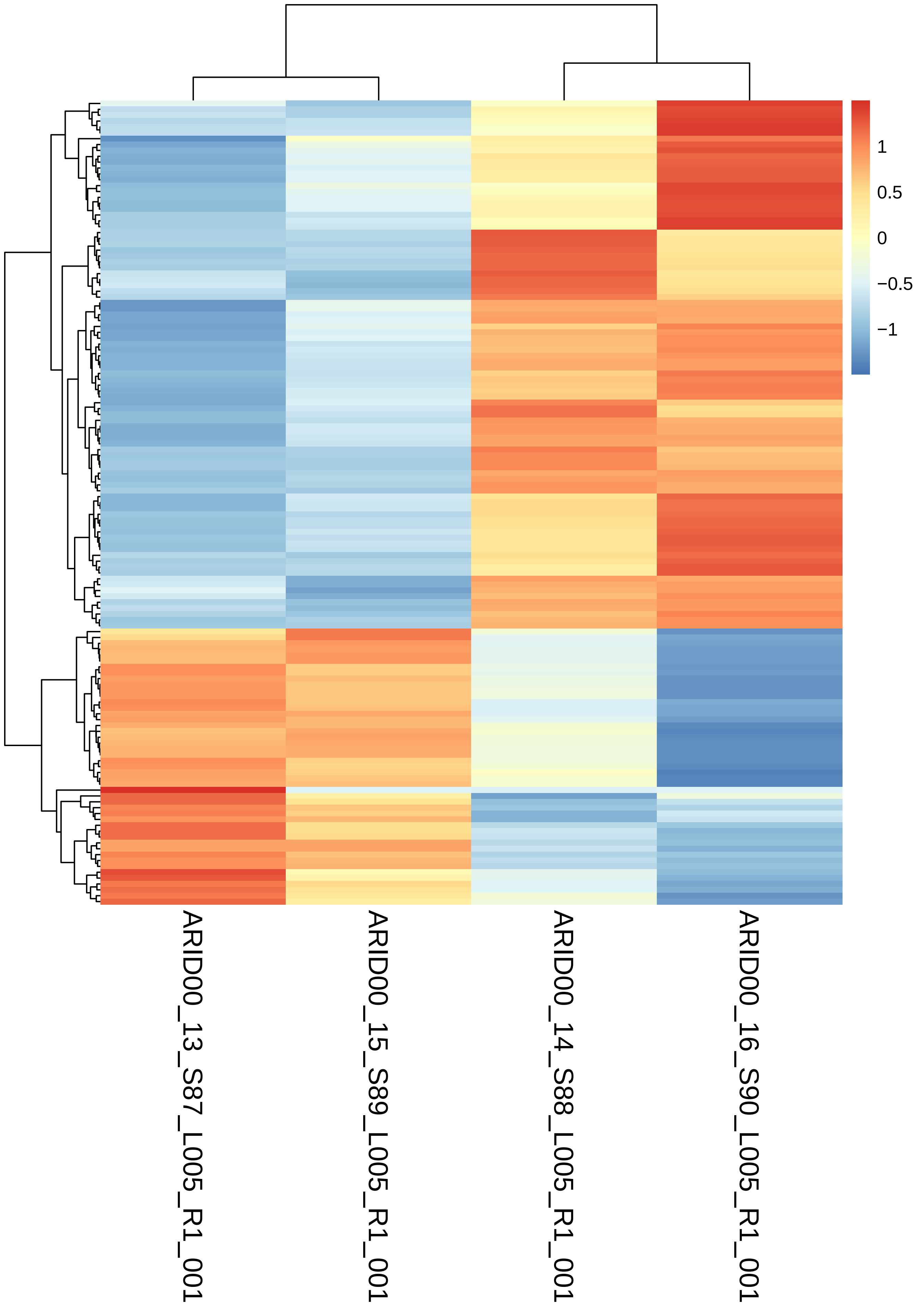
